# Endoribonuclease activity of XRN-2 is critical for RNA metabolism and survival of *Caenorhabditis elegans*

**DOI:** 10.1101/2021.02.25.432855

**Authors:** Moumita Sardar, Aniruddha Samajdar, Adil R Wani, Saibal Chatterjee

## Abstract

microRNAs (miRNAs) are known to regulate a vast majority of the eukaryotic genes by post-transcriptional means, and multiple nucleases play critical roles in the biogenesis and turnover of these regulators. A number of studies have indicated that turnover is important for determining the abundance of miRNAs, and thus, in turn govern their functionality. Recent research in *Caenorhabditis elegans* has revealed an ATP-independent endoribonuclease activity of the ‘miRNase’-XRN-2. Here, we report the characterization of this new enzymatic activity of the fundamentally important XRN-2, and show that it is critical for miRNA turnover and survival of quiescent dauer worms. The dual enzymatic activity of XRN-2 capacitates the mechanism of miRNA turnover to be dynamic, which might confer adaptive advantage to the organism. In continuously growing worms, this new enzymatic activity does not act on miRNAs, but it is important for the generation of mature ribosomal RNAs, which in turn is critical for translation, and thus indispensable for the survival of worms.

---

Ribonucleases play a central role in RNA metabolism. They not only digest and clear out RNAs that have served their purposes, and make way for new RNA-players, but they are also responsible for the processing of innumerable large precursor RNA molecules into mature functional forms (**1, 2**). More than four decades back, 5’-3’ exoribonuclease activity was reported for the first time (**3**), and except one lone bacterial occurrence (**4**), they are found only in eukaryotes. The most widely studied 5’-3’ exoribonucleases belong to the XRN family, which play varied roles in RNA metabolism (**5, 6**). The nuclear localized XRN family member, XRN-2, is known to be critically important for the processing and turnover of a variety of fundamentally important RNAs, including rRNA, snoRNA, TERRA, TSSa, tRNA, and also for the degradation of transcripts emerging from H3K27me3-marked genes (**5, 7**).

In the recent past, XRN-2 and its cytoplasmic paralogue XRN-1 have been reported as ‘miRNases’ in *Caenorhabditis elegans* (*C. elegans*), where they are capable of modulating miRNA function (**8-11**). Very recently, an XRN-2 containing macromolecular miRNA turnover complex (miRNasome-1) has been reported from worms, where XRN-2 has been shown to possess a previously unknown endoribonuclease activity (**12**). ATP binding to the ATP binding/ hydrolysing subunit of miRNasome-1 exerts an inhibitory effect on this endoribonuclease activity. Whereas, miRNA binding to the *bona fide* RNA-binding subunit of miRNasome-1 stimulates ATP hydrolysis, and the energy is utilized towards the transfer of the miRNA to XRN-2 for its exoribonucleolysis. Interestingly, this endoribonuclease activity was shown to be ATP-independent and far more efficacious than the well-known exoribonuclease activity. It was suggested that such an ATP-independent endoribonucleolytic-mode of turnover mechanism would be highly suitable for the worms to withstand a challenging environment, where food is unavailable and thus energy needs to be conserved, and one such physiologically relevant situation might be the miRNA homeostasis during the dauer stage (**12-14**). Continuously growing worms embark on a sturdy alternative quiescent life cycle stage called dauer upon starvation and overcrowding, and continue to live for 3-4 months without feeding, and when the population density reduces and food becomes available, they get back to the continuous life cycle (**13**). Therefore, understanding of the endoribonuclease activity of XRN-2, especially in the context of miRNA turnover and dauer metabolism, might be very important to understand the biology of the quiescent dauer worms, which are analogous to quiescent adult stem cells, and the parasitic nematodes in their infective stages (**13, 15**).

Notably, most of the inferences on XRN-2’s role in RNA metabolism have been drawn using genetic studies, which mostly didn’t involve a biochemical demonstration of the activity on the substrates. Moreover, those studies didn’t exclusively employ mutants of the exoribonuclease activity of XRN-2 (**1, 2, 6, 7**). Thus, it is not clear whether those roles can solely be attributed to the already characterized exoribonuclease activity, and it might be possible that some of those attributed roles are actually carried out by the newly suggested endoribonuclease activity. Therefore, characterization of this enzymatic activity and identification of its endogenous substrates through both *in vivo* and *in vitro* studies would bring about major revelations that would certainly be important to understand the biogenesis and degradation of different species of RNA molecules.

In this study, we report a metal ion-mediated endoribonuclease activity of the fundamentally important enzyme, XRN-2, where the active site gets organized through conformational changes elicited upon substrate (miRNA) binding to a distant domain. This ATP-independent endoribonuclease activity acts on miRNAs in energy-deprived quiescent dauer worms, and its perturbation severely affects miRNA homeostasis, which in turn affects the target mRNAs, and the worms succumb. Conversely, this activity doesn’t affect miRNAs in continuously growing worms, where the ATP-dependent exoribonuclease activity of XRN-2 appears to play the role. However, when this exoribonuclease activity is not allowed to access energy through depletion of the ATP hydrolyzing subunit of miRNasome-1, the ATP-independent endoribonuclease activity of XRN-2 bails out the situation by maintaining mature miRNA levels. Thus, this previously uncharacterized nuclease activity incorporates a layer of dynamism to the mechanism of miRNA turnover. Lastly, we demonstrate that the endoribonuclease activity of XRN-2 is critical for RNA metabolism beyond miRNAs, as it is involved in the processing of precursor ribosomal RNAs, and thus, very important for translation.

## RESULTS

### XRN-2 is critical for miRNA metabolism and survival of quiescent dauer worms

XRN-2 is an important ‘miRNase’ in the worms and critical for its miRNA metabolism. Recently, an XRN-2-containing miRNA turnover complex, miRNasome-1, was reported to be capable of acting in two alternative modes. An ATP-independent endoribonucleolytic mode of action was shown to be more efficacious than its ATP-dependent exoribonucleolytic mode of action. Since, continuously growing worms would have access to food and energy, this ATP-independent mode of functioning might be useful, when food is scarce and energy needs to be conserved. Thus, it was implicated for miRNA homeostasis during the quiescent dauer stage (**12, 13**). Indeed, the relative cellular ATP levels in 4-week old dauer worms are heavily diminished (∼10 folds) compared to that of the continuous life cycle stage worms (L2, L3 and L4, **Fig. 1A**). Consistent with previous observations (**16, 17**), we noticed that the levels of some of the potential miRNasome-1 substrate miRNAs in the dauer stage are diminished than that observed during the relevant normal life cycle stages (L2, L3, **Fig. 1B**). We further observed that the precursor levels of those highly diminished mature miRNAs were not changed, and thus, indicating it to be an event happening at the level of the mature miRNA only (**Fig. 1C, data not shown**). Importantly, we could find that perturbation of XRN-2 (see methods) led to the accumulation of several mature miRNAs in dauer worms, without affecting the levels of their precursors (**Fig. 1D, data not shown**), which in turn down-regulated target mRNAs known to be important for dauer metabolism (**Fig. 1E)**. This clearly suggested that XRN-2 specifically acts on mature functional miRNAs, and thus, it is a *bona fide* ‘miRNase’ even in the quiescent dauer stage. Notably, those experimental dauer worms succumbed within 100 hours, whereas the control dauers showed very little mortality and continued to live beyond 50 days (**Fig. 1F**).

**Fig 1.**
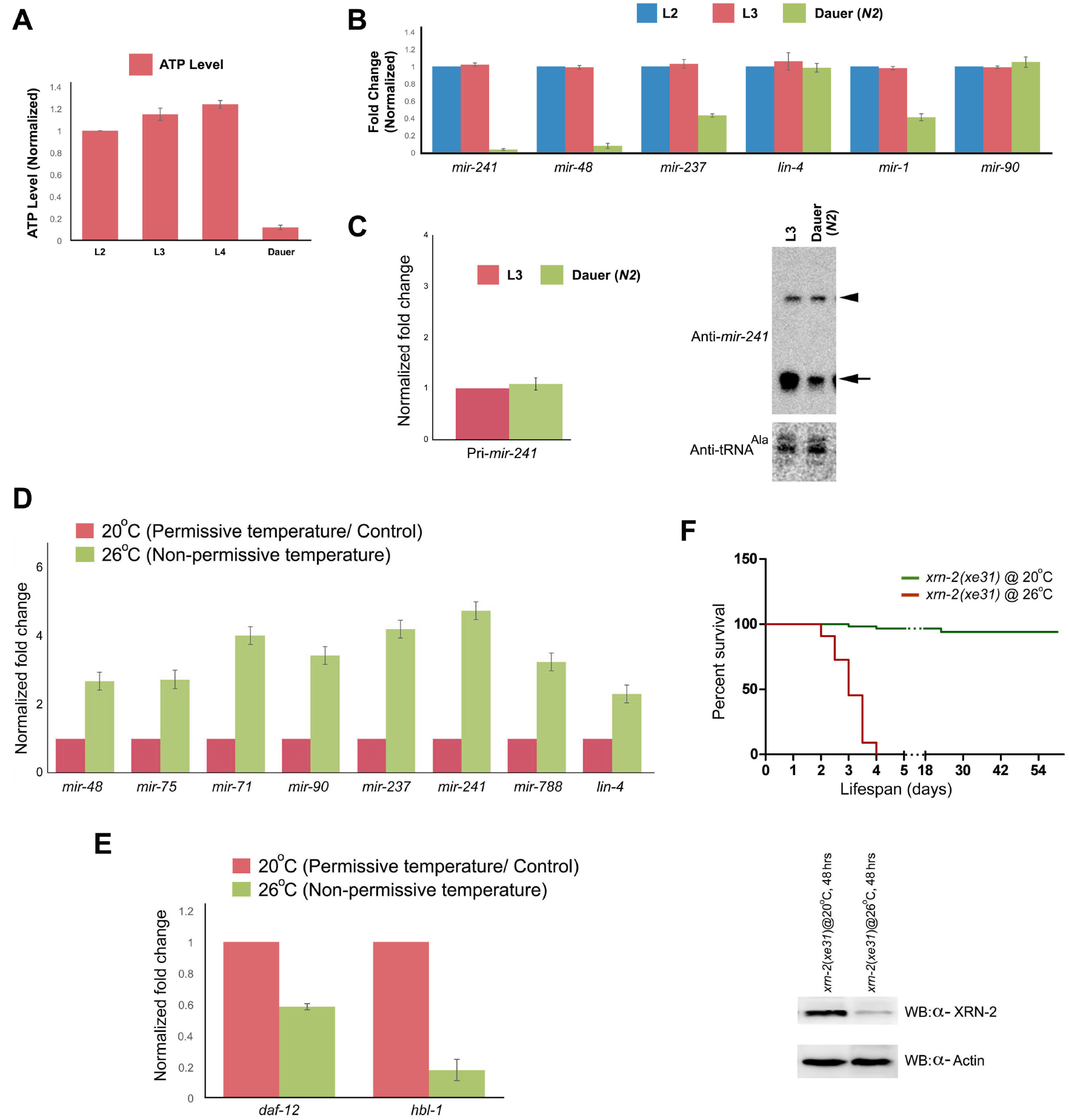
XRN-2 is crucial for miRNA metabolism and survival of dauer worms. **(A)** Diminished ATP levels in dauer worms compared to normal life cycle stage (L2, L3, and L4) worms. **(B)** Some miRNA levels are significantly reduced in dauer worms compared to the worms from equivalent normal life cycle stages (L2, L3), when pri-miRNA (eg. pri-*mir-241*, **C left panel**) and pre-miRNA (eg. pre-*mir-241*, indicated with arrowhead, **C right panel**) levels in these two groups remain unchanged. **(D)** A number of mature miRNAs, including some key *let-7* family members, are accumulated in *xrn-2*(*xe31*) mutant dauer worms, upon depletion of XRN-2 (error bars indicate SEM of at least three biological replicates). Unlike continuously growing worms, *lin-4* shows accumulation upon XRN-2 perturbation in dauer worms (reference 12). **(E)** Levels of *let-7* and *lin-4* family targets, *hbl-1* and *daf-12*, crucial for dauer metabolism, are diminished upon XRN-2 depletion (error bars indicate SEM of at least three biological replicates) in *xrn-2*(*xe31*) mutant dauer worms. **(F) Top panel**. Dauer worms of the temperature sensitive *xrn-2*(*xe31*) mutant succumb, when they are maintained at the non-permissive temperature (26°C). **Bottom panel**. XRN-2 is depleted in *xrn-2*(*xe31*) mutant dauer worms maintained at the non-permissive temperature (for 48 hrs).

### XRN-2 employs a metal ion-mediated catalytic mechanism for its endoribonuclease activity

The above results clearly suggested that XRN-2’s ATP-independent endoribonuclease activity holds potential to shape the miRNA landscape in dauers (**Fig. 1**), which in turn might play a crucial role in the adaptation and survival of worms. Thus, we decided to characterize this enzymatic activity. XRN proteins are known to be distantly related to flap endonuclease FEN-1 and becteriophage T4 RNase H, which are magnesium-dependent nucleases and members of the PIN domain family (**Fig. 2A, reference 5**). Bioinformatic analyses with the available atomic structure of worm XRN-2 (5FIR.pdb, **reference 18**) identified amino acid residues in XRN-2, which are homologous to two out of the three critical catalytic core residues of human FEN1 endoribonuclease, and three out of the three critical catalytic core residues of T4 RNase H (**Fig. 2B, C;** 3Q8L.pdb, 1TFR.pdb; **references 19-21**). Of note, two out of the three catalytic core residues of T4 RNase H are homologous to the active site residues of FEN-1 endoribonuclease. Interestingly, the homologous residues in XRN-2 are located towards the dead-end of XRN-2’s exoribonuclease active site tunnel, beyond the residues necessary for exoribonucleolysis (**Fig. 2D**). Therefore, in this arrangement, two activities can never act at the same time, and would necessitate the protein to be in two alternative states, where a given state would be capable of carrying out only one of the two activities. But, mutation of those critical residues in XRN-2 didn’t affect its endoribonuclease activity at all (**Fig. 2E**). These results suggested that XRN-2 might be hosting a novel endoribonuclease active site, and possibly a catalytic mechanism, not known before, and to unravel that we decided to adopt a more generalized and systematic approach.

**Fig 2.**
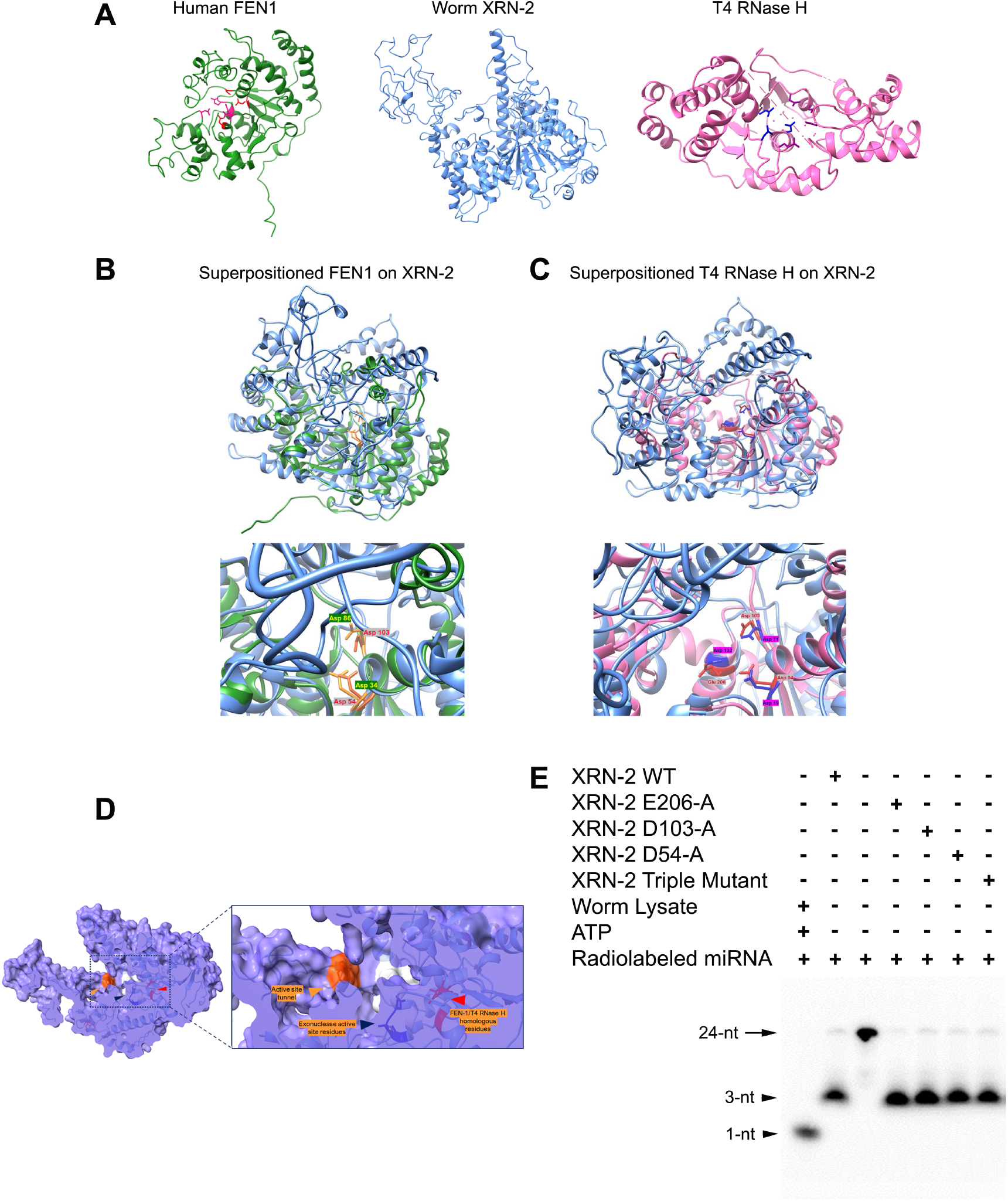
XRN-2 residues homologous to catalytic core residues of T4 RNase H and FEN-1 endoribonuclease are present deep inside its exoribonuclease active site tunnel. **(A)** Atomic structure of human FEN1 (left panel), *C. elegans* XRN-2 (centre), and T4 RNase H (right panel). **(B and C) Top panels**. Comparison of *C. elegans* XRN-2 (Blue) with human FEN-1 (Green) and T4 RNase H (Pink) by superposition. **Bottompanels**. Close-up view showing the positioning of the homologous residues in the super positioned proteins. **(D)** A cut-section of the space-filling model of XRN-2 reveals the position of its residues homologous to T4 RNase H and FEN-1 endoribonuclease active site residues (indicated with red arrowhead), towards the dead-end of the exoribonuclease active site tunnel, beyond the residues necessary for exoribonucleolysis (indicated with blue arrowhead). A close-up view is provided on the right-hand side. The bulky amino acid (W, in orange) at the point of entry to the active site tunnel is indicated with an orange arrowhead. **(E) XRN-2 residues homologous to T4 RNase H and FEN-1 endoribonuclease active site do not contribute to its endoribonuclease activity**. miRNA turnover assay performed with the purified recombinant wild type, single mutant, and triple mutant (D54-A, D103-A, E206-A) XRN-2 proteins, as indicated. Mutations fail to inhibit the endoribonuclease activity of XRN-2. Two out of the three catalytic core residues of T4 RNase H are homologous to two out of the three active site residues of FEN-1 endoribonuclease. D103 and D54 in XRN-2 are homologous to those two critical residues.

Employing several truncated versions of the recombinant XRN-2, we could localize the endoribonuclease activity to its N-terminal fragment (**Fig. 3A**). This 474 amino acid (aa) long fragment houses a number of Histidine residues, which could be involved in an acid-base catalytic mechanism to achieve the endoribonucleolytic cleavage (**Fig. 3B right panel, reference 1**). Conversely, a large number of Aspartic acid and Glutamic acid residues present in this fragment could also be responsible for liganding metal ions, which in turn would coordinate hydroxyl groups to perform nucleophilic attack on the substrate phosphodiester backbone to achieve endoribonucleolytic cleavage (**Fig. 3B left panel, reference 1**). In order to identify the underlying mechanism (acid-base catalysis vs metal ion-mediated catalysis), we performed an *in vitro* miRNA turnover assay involving a 5’-radiolabeled miRNA and recombinant XRN-2, where the 3-nt long endoribonucleolytic products were further subjected to 5’-radiolabeling using T4 Polynucleotide Kinase (PNK, **Fig. 3C**). Any increase in the signal for the 3-nt long product would indicate that the endoribonucleolytic mechanism involved was acid-base catalysis, which results in the formation of 5’-OH products, and thus allows subsequent 5’-phosphorylation. Contrastingly, no increase in signal for those 3-nt long products would reveal a metal ion-mediated mechanism that produces products with 5’-phosphates, which can’t be further phosphorylated (**Fig. 3C)**. The assay results showed the latter to be true for XRN-2’s endoribonuclease activity **(Fig. 3C, D)**. This result was reconfirmed when we obtained a radiolabeled monoribonucleotide, alongside 3-nt long products, upon incubation of a 22-nt long body-radiolabeled *let-7* (where the 22^nd^ base is U) with recombinant XRN-2, which cleaves the substrate after every three nucleotides (data not shown, **reference 12)**. Only a metal ion-mediated mechanism would cleave the bond between the 21^st^ and 22^nd^ nucleosides to give rise to a nucleotide with a 5’-radiolabeled alpha phosphate. Whereas, an acid-base catalytic mechanism would have generated a nucleoside with a 5’-hydroxyl end, and thus, it would have remained undetected in the autoradiogram.

**Fig 3.**
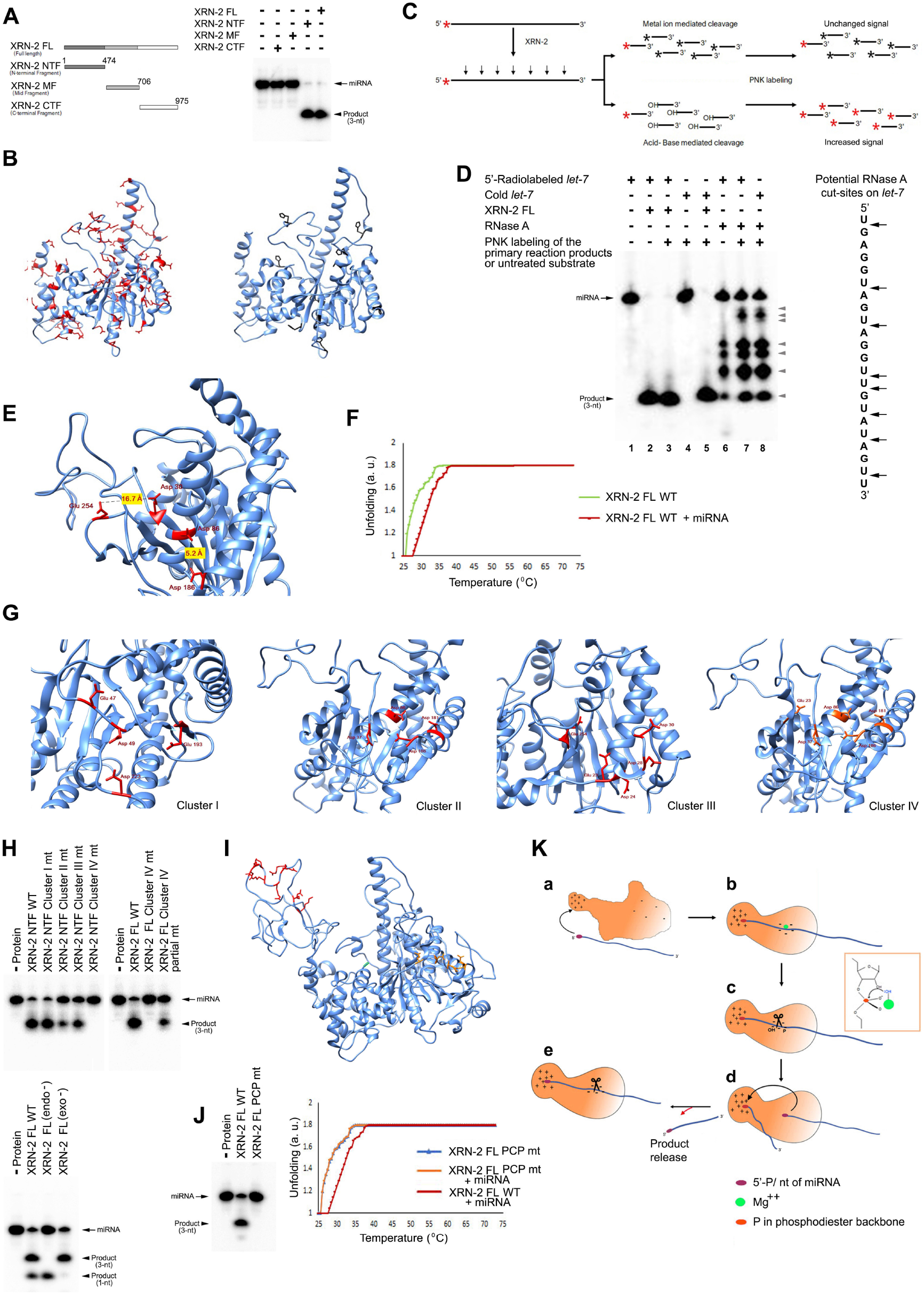
XRN-2 adopts a metal ion-mediated catalytic mechanism for its endoribonuclease activity. **(A) Left panel**. Schematic of the full length, and the different truncated versions of the recombinant XRN-2 protein. **Right panel**. The N-terminal fragment (NTF) of XRN-2 harbors the highly processive endoribonuclease activity. 10 nM body-labeled *mir-84* was incubated with 9 nM of recombinant proteins. **(B) Left panel**. Ribbon representation of the worm XRN-2 NTF protein structure (5FIR.pdb) with highlighted Aspartic acid and Glutamic acid residues (stick representation in red) capable of organizing a metal ion-mediated catalysis to achieve endoribonucleolysis. **Right panel**. Histidine residues highlighted in black (stick representation) capable of organizing an acid-base catalytic mechanism. **(C)** Schematic of an assay to distinguish between a metal ion-mediated and acid-base catalytic mechanism employed by XRN-2 to achieve endoribonucleolysis of substrate miRNA. **(D) Left panel**. Unchanged 3-nt long endoribonucleolytic product upon PNK treatment reveals a metal ion-mediated catalytic mechanism employed by recombinant XRN-2 protein (compare lanes 2 and 3). RNase A serves as a positive control for acid-base catalytic activity, whose 5’-OH products got further radiolabeled by PNK (compare lanes 6-8, grey arrowheads). Lanes 4 and 5 show robust 5’-PNK labeling of 5’-hydroxylated miRNA at a concentration same as the employed substrate in the experimental lanes, and its further robust degradation by recombinant XRN-2, respectively. **Right panel**. The potential RNase A cleavage sites on the substrate miRNA used in the assay (*let-7*). **(E)** N-terminal region of the crystal structure of XRN-2 depicting the two closest (D86, D186) and farthest (D30, E254) residues that are 5.2 and 16.7 Å apart, respectively, which may ligand metal ions. **(F)** Melting assay demonstrates the requirement of a higher temperature for unfolding of recombinant full length XRN-2 in the presence of miRNA. **(G)** N-terminal regions of the crystal structure of XRN-2 depicting four different sets of residues (clusters I-IV) having potential to ligand metal ion required for metal ion-mediated catalysis. **(H) Top panel**. Mutation of the residues in cluster IV abrogates the endoribonuclease activity of XRN-2. **Bottom panel**. Abrogation of the endoribonuclease activity of XRN-2 doesn’t affect its exoribonuclease activity. ∼90 nM recombinant XRN-2 and high substrate concentration (∼110 nM) was used to achieve the exoribonucleolytic product (monoribonucleotides). **(I)** Ribbon representation of the worm XRN-2 protein structure (5FIR.pdb) with highlighted residues (stick representation in red) forming a potential positively charged patch (PCP) for binding RNA. Catalytic core residues are shown in brown. **(J) Left panel**. Mutation of the residues in the PCP abolishes the endoribonuclease activity of XRN-2. **Right panel**. PCP mutant XRN-2 fails to show any change in melting pattern in the presence of miRNA. **(K) A schematic model for the endoribonuclease activity of XRN-2**. miRNA binding-induced conformational changes are indicated through alteration in the shape of the XRN-2 cartoon, and changes in the proximity of catalytic core residues indicated with negative signs (b). Inset depicts a liganded metal ion coordinating a hydroxyl group, which in turn performs a nucleophilic attack on the phosphodiester bond of the RNA molecule.

### Endoribonuclease active site of XRN-2 gets organized through conformational changes elicited upon substrate miRNA binding to a distant domain

Employing SWISS-MODEL, we generated a XRN-2 structure by ribbon overlaying its entire sequence on the available atomic structure of worm XRN-2, which was derived from a protein with some deletions (5FIR.pdb, **reference 18**). It revealed a number of potential metal ion liganding residues in the aforementioned N-terminal fragment, but none of those appear to be close enough (3-4 Å) to form a pair, as minimally required to ligand a metal ion. Indeed, the two closest potential residues (D86 and D186) residing in a domain are 5.2 Å apart, and the farthest are 16.7 Å apart (D30 and E254, **Fig. 3E**). Interestingly, a thermal shift assay (**18**) with the recombinant full length XRN-2, as well as the truncated 5’-fragment (474 aa), in the presence of substrate miRNA demonstrated the requirement of a much higher temperature to unfold the protein for dye binding (3.9 °C **±** 0.35; **Fig. 3F**, data not shown), compared to the assay performed without miRNA. This result suggested that miRNA binding increases stability of XRN-2 by inducing conformational changes, and that in turn might bring the critical residues close enough to form a metal ion-mediated catalytic site, which are otherwise located far from each other, as evident in the crystal structure. We could identify three locations on the active 5’-fragment of XRN-2, where four/ five critical residues are concentrated maximally ∼14 Å apart from each other to form clusters, and thus might be suitable for metal ion-mediated catalysis (**Fig. 3G**). Notably, four/ five negatively charged residues are likely to ligand metal ions strong enough to result in a highly efficacious endoribonuclease activity, compared to that of the intrinsic weak exoribonuclease activity of XRN-2 carried out through only two metal ion liganding residues (D234, D236), as evident through *in vitro* turnover assays reported here and before (**Fig. 3H bottom panel, reference 12**). When all the residues in cluster III were mutated, it showed a modest reduction in the *in vitro* endoribonuclease activity of XRN-2. But, a substantial reduction (79% + 2%) in the activity was observed upon mutation of cluster II residues, and each of the residues of this cluster was found to be important for the activity (**Fig. 3H top panel**, data not shown). Further mutational analyses revealed that mutation of only a single residue (E23) of cluster III is responsible for the observed reduction in the endoribonuclease activity, whereas, mutation of the other residues of this cluster has no effect (data not shown). E23 happens to be very close to cluster II (**Supplementary figure 1**), and thus, we added E23 to cluster II to create a new cluster IV (**Fig. 3G right-most panel**). Indeed, mutation of all the residues of cluster IV (E23, D37, D86, D181, D186) resulted in a complete abrogation of the *in vitro* endoribonuclease activity of XRN-2. Therefore, they essentially form the catalytic core to ligand the metal ion, although, we do not know how many of those are required for the optimal activity (**Fig. 3H top panel**). Notably, a widely used bio-informatics tool (MIB: Metal ion binding site prediction and docking sites, **reference 22**) predicted that D37 and D86, which are constituents of this biochemically defined endoribonuclease active site, are suitable for docking divalent magnesium. The endoribonuclease activity diminished to about 50%, when two specific residues (D86, D181) of the catalytic core were mutated (**Fig. 3H top panel, right-most lane**). Importantly, perturbation of the endoribonuclease activity didn’t affect the *in vitro* exoribonuclease activity of XRN-2 (**Fig. 3H bottom panel**), which could be achieved at high protein and substrate concentrations, as reported before (**12**).

Interestingly, a thermal shift assay with the recombinant wild type and endoribonuclease mutant XRN-2 proteins, in the presence of substrate miRNA demonstrated a requirement of a much higher temperature to unfold for dye binding, compared to that of the wild type protein alone (**Supplementary figure 2A**). Notably, in the presence of miRNA, the wild type protein required modestly higher temperature for melting than the mutant protein. The above result suggested that the catalytic core residues might not play any primary role for binding the substrate miRNA. Therefore, we looked for surface localized potential RNA-interacting residues (mostly positively charged residues) in close proximity with each other, which might act as a binding patch for a negatively charged RNA and elicit a conformational change to optimally re-position the catalytic core residues to ligand metal ions. We found two such clusters of surface localized residues, which were less than 11.5 Å away from the nearest catalytic core residue (**Supplementary figure 3A, B**). But, their mutation had no effect on the *in vitro* endoribonuclease activity of XRN-2 (**Supplementary figure 3C**). Interestingly, we observed another cluster of nine closely placed potential RNA-interacting residues forming a **p**ositively **c**harged **p**atch (PCP) on the surface of the protein (K421, R423, R432, R435, R437, R439, R441, Q443, Q444; **Fig. 3I, Supplementary figure 3B)**, but located as far as 47.7 Å away from the nearest catalytic core residue (E23, **Fig. 3G right-most panel, I, K**). A recombinant protein mutant for these PCP residues showed no endoribonuclease activity (**Fig. 3J left panel)**. The distance between this PCP and the catalytic core, which may be the length of around three nucleotides upon conformational reorientations, might act as a molecular ruler and facilitate the formation of three nucleotide long products (**Fig. 3I, K**). However, considering the available atomic structure of worm XRN-2 (5FIR.pdb), the PCP residues are far (47.7 Å) away from the catalytic core, which suggests that this PCP must get close to the active site residues through conformational changes upon binding the substrate miRNA. Of note, the protein mutant for the PCP residues was unable to show any temperature shift in melting assays in the presence of miRNA (**Fig. 3J right panel**), and that again indicated that substrate miRNA binding indeed elicits conformational changes in XRN-2 suitable to carry out endoribonucleolysis of the substrate. Additionally, Circular Dichroism (CD) spectroscopic analyses of the recombinant mutant proteins showed no changes in their secondary structures (data not shown), and thus confirmed that the loss of functions are not due to non-specific reasons.

### The endoribonuclease activity of XRN-2 is responsible for miRNA turnover in energy-deprived dauer worms, but not the ones in continuous life cycle

A homozygous null mutant worm for the endoribonuclease activity of XRN-2 proved to be lethal, but a homozygous worm with mutated two of the catalytic core residues (D86-A, D181-A) could be generated. This mutant worm strain *xrn-2*(*PHX25*) grew by and large normally at 20°C, but at the non-permissive temperature (26°C), it showed a number of phenotypes and eventually arrested and died in the ninth generation (F9, **Supplementary Table 1, Supplementary figure 4A**). Some F9 worms turned L4, but the development didn’t proceed any further. We chose F4 worms for molecular analyses, since, strong phenotypes like larval arrest was first observed in this generation, alongside other phenotypic defects. Late generations were also avoided as they might accumulate different secondary effects. The RNA from the L4 stage of the F4 worms growing at 26°C was compared with that of the worms growing at 20°C, at the same generation and stage. We found no significant change in the levels of their endogenous mature miRNAs (**Supplementary figure 4C**), which indicated that this activity might be important for other substrate RNAs in these continuously growing worms. Accordingly, the lysate from these F4 experimental worms showed no change in the XRN-2-dependent and ATP-driven exoribonuclease activity on radiolabeled substrate miRNA in an *in vitro* turnover assay (**Supplementary figure 4D**). This suggested that in the experimental worms, the exoribonuclease activity of the endogenous XRN-2 was unaffected. We also found no change in the levels of XRN-2 protein in these samples **(Supplementary figure 4B)**, which confirmed that the mutations do not exert any non-specific effect on XRN-2 stability.

Contrastingly, the dauer worms of this mutant strain [*xrn-2*(*PHX25*)] maintained at the non-permissive 26°C, as opposed to the ones maintained at the permissive 20°C, showed considerable accumulation of several mature miRNAs without any significant change in the levels of their precursors (**Fig. 4B-D**). We had included certain *let-7* and *lin-4* family members (*mir-241, lin-4* etc) in the cohort of miRNAs we examined, as their mature forms were known to be present at diminished levels in dauers compared to the equivalent stages in continuously growing worms, and were already shown to regulate specific mRNA targets critical for dauer metabolism (**16, 17**). Thus, the above results clearly suggested that the endoribonuclease activity of XRN-2 is critical for the establishment and maintenance of diminished levels of those miRNAs in dauers, and it acts only on the mature forms. The accumulated mature miRNAs in dauer worms maintained at 26°C were active as we could find that some of the validated targets (eg. *daf-12, hbl-1*) of *let-7* and *lin-4* family members were diminished compared to the sample from the dauers maintained at 20°C (**Fig. 4E**). The XRN-2 protein level in these experimental dauers remained comparable to that of the control dauers (**Fig. 4A bottom panel**), which suggested that the specific mutations in XRN-2’s endoribonuclease active site have indeed resulted in a deregulation of the endogenous mature miRNA levels in the experimental dauer worms. Importantly, those experimental dauer worms growing at the non-permissive temperature succumbed within 10 days, whereas the dauers at the permissive temperature exhibited very little mortality and continued to live beyond 50 days (**Fig. 4A**). Notably, dauers generated from wild type N2 worms showed similar survival at both the permissive and non-permissive temperatures, which excluded any possible effect of temperature on the lifespan (data not shown). These results clearly indicated that the endoribonuclease activity of XRN-2 is critical for miRNA turnover and survival of energy-deprived dauer worms, but does not affect the miRNA metabolism of worms during their continuous life cycle. This further implied that only the exoribonuclease activity of XRN-2 is responsible for miRNA turnover during the continuous life cycle.

**Fig 4.**
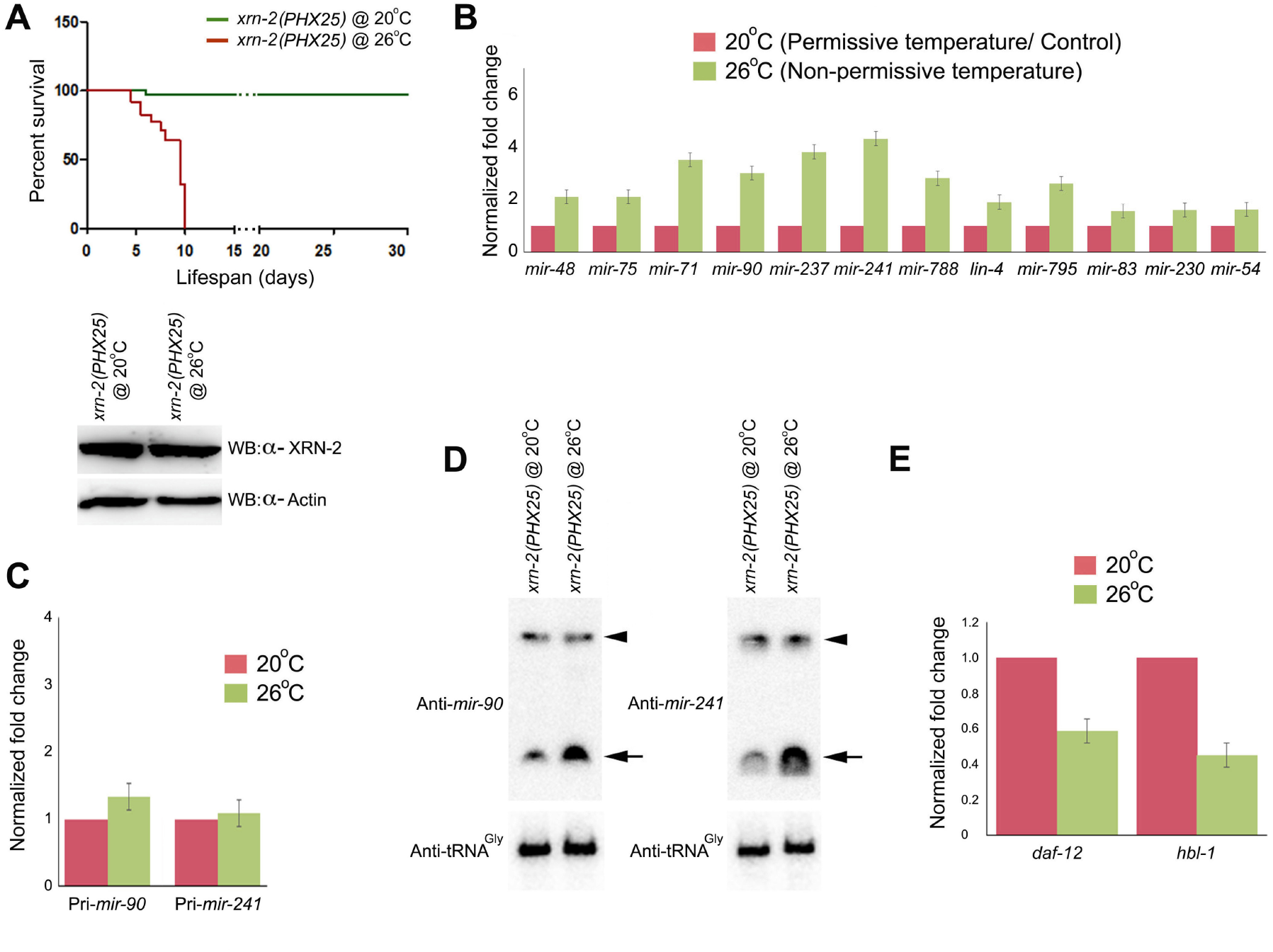
Endoribonuclease activity of XRN-2 is critically important for miRNA metabolism and survival of dauer worms. **(A) Top panel**. Dauer worms of the temperature sensitive *xrn-2*(*PHX25*) strain having mutations in the endoribonuclease active site succumb, when they are subjected to non-permissive temperature (26°C). The first death was recorded on the fifth day, and this time point was selected for molecular analyses. **Bottom panel**. Stability **of** XRN-2 protein remain unchanged in *xrn-2*(*PHX25*) dauer worms maintained at the non-permissive temperature (for ∼ 110 hrs). **(B)** A number of mature miRNAs, including some key *let-7* family members, accumulate in *xrn-2*(*PHX25*) dauer worms under the experimental conditions (error bars indicate SEM of at least three biological replicates). Unlike continuously growing worms, *lin-4* shows accumulation in dauer worms (reference 12). **(C, D)** RT-qPCR (**C**) and northern blot (**D**) analyses demonstrate no significant changes in the levels of pri-miRNAs and pre-miRNAs (indicated with arrowheads), respectively. **(E)** Levels of *let-7* and *lin-4* family targets, *daf-12* and *hbl-1*, crucial for dauer metabolism, get diminished in *xrn-2*(*PHX25*) dauer worms subjected to non-permissive temperature (error bars indicate SEM of at least three biological replicates).

### Endoribonuclease activity of XRN-2 confers dynamism to the mechanism of miRNA turnover in worms

It has already been demonstrated that at optimal miRNA concentration, the miRNasome-1-residing XRN-2 can act in an ATP-dependent exoribonucleolytic mode, when miRNasome-1.4 supplies energy for that action by hydrolysing ATP. miRNasome-1.4 also inhibits the endoribonuclease activity of XRN-2 upon ATP binding. But, depletion of miRNasome-1.4 doesn’t lead to any accumulation of miRNAs in continuously growing worms (**reference 12, Supplementary figure 5 top panel**). It is most likely that if miRNasome-1.4 would be absent or inhibited, the miRNasome-1-residing XRN-2 would still be able to act endoribonucleolytically on the substrate miRNA, which would be sufficient to inactivate the miRNA, and thus, will not allow any accumulation of mature miRNAs. Accordingly, if miRNasome-1.4 would be depleted by RNAi in *xrn-2*(*PHX25*) worms at 26°C, that might limit the endoribonuclease activity mediated compensation for the missing exoribonuclease activity, and would in turn affect the levels of mature miRNAs. Indeed, it resulted in accumulation of a number of mature miRNAs (**Supplementary figure 5 bottom panel**), which didn’t change upon depletion of miRNasome-1.4 alone in the wild type worms. The above result indicated that the endoribonuclease activity of XRN-2 enables the miRNasome-1-mediated miRNA turnover mechanism to be dynamic in response to availability of energy in the environment. Thus, together, our data suggest that a bimodal mechanism for turnover of mature miRNAs is critical for miRNA metabolism in *C. elegans*, and its adaptation.

### Endoribonuclease activity of XRN-2 takes part in ribosomal RNA metabolism in worms

The most striking features of the continuously growing *xrn-2*(*PHX25*) worms mutant for the endoribonuclease activity, maintained at the non-permissive temperature (26°C), were reduction in the number of embryos formed, reduced hatching of embryos into L1 worms, and subsequent growth arrest at several larval stages (**Supplementary figure 4A, Supplementary Table 1**). It is well known that ribosome biosynthesis is critical for germline stem cell differentiation, propagation and maturation of germ cells, and its perturbation can affect gonadogenesis, and also result in postembryonic growth arrest (**23-25**). Notably, XRN-2 has been implicated in the biogenesis and surveillance of ribosomal RNAs (rRNAs), where it facilitates the formation of mature 5.8s and 28s rRNAs by removing precursor sequences (**26-29**). Therefore, we decided to check the status of the different rRNA species in those continuously growing worms mutant for the endoribonuclease activity of XRN-2 at the non-permissive temperature (**Fig. 5A**). Again, for our analyses, F4 worms were chosen for the reasons described before. We observed accumulation of additional bands larger than the mature 5.8s and 28s rRNAs, with a concomitant decrease in the levels of the mature forms in the experimental samples. However, the mature 18s rRNA production remained unaffected, and that excluded the possibility of any transcriptional aberrations contributing to the depletion of mature 5.8s and 28s rRNAs. Of note, a distinct 5.8s rRNA precursor band could also be detected in the *xrn-2*(*PHX25*) worms at the permissive temperature, but not in the wild type *N2* worms grown at either of the temperatures. Thus, even in the permissive temperature, the mutation affected the biogenesis of mature 5.8s rRNA, although, that didn’t prove to be sufficient to exert a significant effect on the worm development (**Supplementary figure 4A, Supplementary Table 1**). Additionally, XRN-2 has also been shown to be important for the biogenesis of snoRNAs (**5, 28**), and we observed changes in the abundance of certain snoRNA species in the *xrn-2*(*PHX25*) worms grown at the non-permissive temperature (data not shown).

**Fig 5.**
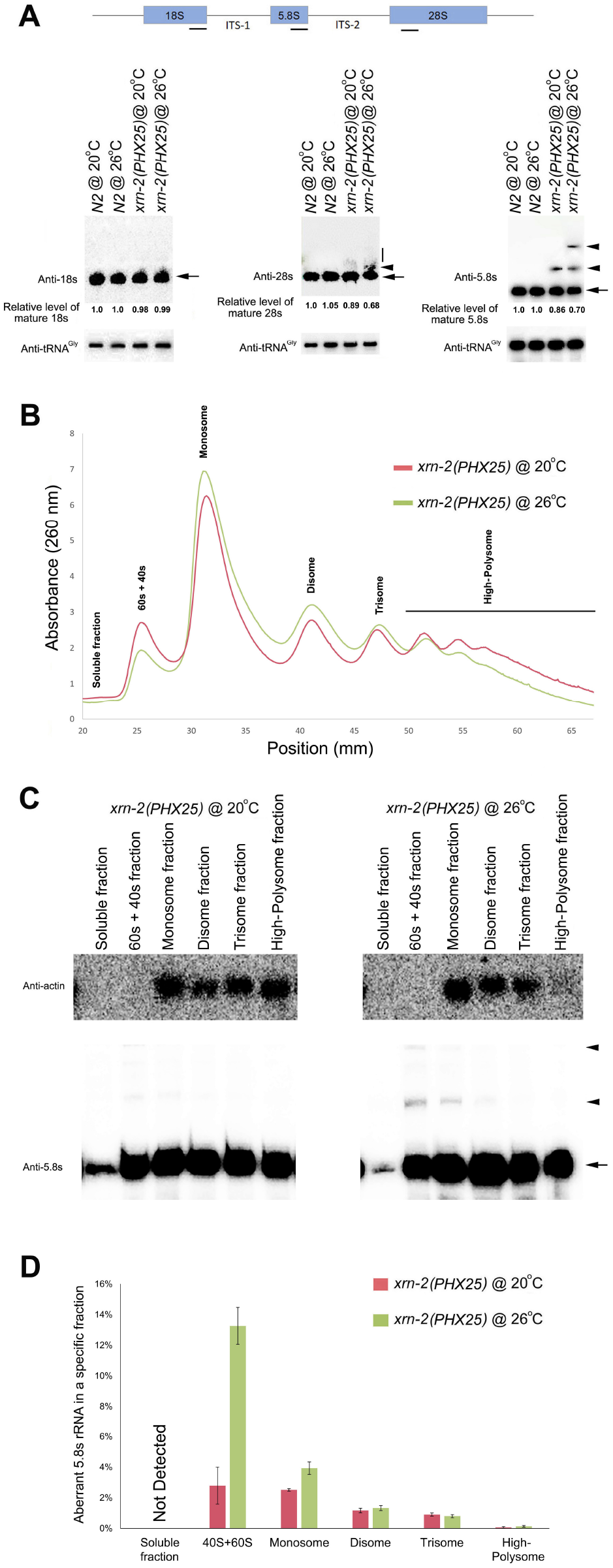
Perturbation of the endoribonuclease activity of XRN-2 causes defects in ribosomal RNA processing. **(A) Top panel**. Schematic diagram depicting the 35s pre-rRNA, where the underlying small bars indicate to the approximate positions to which the northern probes are directed for detecting mature 18s, 5.8s, and 28s rRNAs. **Bottom panel**. Northern blot analyses reveal depletion of mature 5.8s and 28s rRNA, as well as accumulation of aberrant precursor bands, in total RNA samples extracted from *xrn-2(PHX25)* worms in their continuous life cycle, grown at the non-permissive temperature of 26°C. Precursors are also detected in the mutant worms at the permissive temperature, but not in the wild type *N2* worms. Mature 18S rRNA levels remain largely unchanged in all the examined samples. Mature rRNA bands have been indicated with arrows, aberrant precursors with arrowheads, and a vertical bar represents smearish aberrant precursors of 28s rRNA in the mutant worm samples. The samples were resolved on a 1.5% Formaldehyde Agarose denaturing gel and subjected to northern probing **(B)** Polysome-profiles of the two indicated samples exhibit distinct patterns. The different fractions/ peaks have been annotated. The fraction containing the ribosomal subunits is depicted as 40s + 60s. Under our conditions they did not segregate to form distinct peaks. The fractions containing mRNAs loaded with more than three ribosomes have been collectively referred as ‘high-polysome’. **(C) Top panel**. Actin mRNA is abundant in the high-polysome fraction of the sample prepared from *xrn-2(PHX25)* worms grown at permissive 20°C, but largely absent in the same fraction of the experimental sample [*xrn-2(PHX25)* @ 26°C]. The samples were resolved on a 1.5% Formaldehyde Agarose denaturing gel and subjected to northern probing. All lanes are from the same blot. **Bottom panel**. Aberrant precursor forms (indicated by arrowheads) of 5.8s rRNA are well detected in the 40s + 60s, monosome and disome fractions of the experimental sample, but remain undetected in the high-polysome fraction of the same sample. High-polysome fraction of the experimental sample depicts reduction in the mature 5.8s rRNA. The samples were resolved on a 6% Urea PAGE and subjected to northern probing. All lanes are from the same blot. **(D)** Percentage of aberrant 5.8s rRNA precursors in different polysome-profile fractions of the indicated samples. Depicted data represent means ± SEM (n = 3).

We observed that some of the accumulated aberrant 28s rRNA bands were weak and smearish in nature (**Fig. 5A**). This could be due to their inherent instability or rapid degradation as they might have adverse effects on cellular processes like translation. Accordingly, we decided to perform polysome-profiling using the aforementioned experimental worm samples [*xrn-2*(*PHX25*) @ 26°C], and study the effect on translation, if any. We observed a significant diminishment in high-polysome fraction (**Fig. 5B**), compared to the control profile [*xrn-2*(*PHX25*) @ 20°C], indicating reduced translation. This indication was strengthened as we observed a more abundant localization of the highly translated actin mRNA in the monosome fraction, rather than in the polysomal fractions as seen in the control polysome-profile (**Fig. 5C top panel**). Also, the experimental samples presented significant increase in monosome, disome, and trisome fractions. Conversely, the fraction with ribosomal subunits (40s + 60s) showed a substantial decrease in the experimental samples (**Fig. 5B**). This suggested that ribosomal subunits are under-produced, and might also be defective in nature, which is why they failed to progressively load on to mRNAs to form functional high-polysomes to drive translation.

RNA was extracted from the abovementioned polysome-fractions and probed for the different rRNAs. Here, we have focused our analyses on 5.8s rRNA. In the experimental samples, we observed the presence of the aberrant 5.8s rRNA precursors in the fractions containing the ribosomal subunits (40s + 60s), monosome and the disome, whereas, the high-polysome fraction was devoid of the aberrant precursors (**Fig. 5C bottom right panel, 5D**). Notably, the percentage of these aberrant 5.8s rRNAs in the fraction containing ribosomal subunits (40s + 60s) of the experimental samples was significantly higher than that observed in the same fraction of the control samples (**Fig. 5D**). These results supported the aforementioned suggestion that accumulation of aberrant precursor rRNA in the experimental samples increased the population of defective ribosomal subunits, and also ribosomes, which were probably compromised for translation, and did not facilitate formation of high-polysome to drive protein synthesis. Reduced abundance of mature 5.8s rRNA in the 40s + 60s fraction of the experimental samples might also have contributed to the reduction in total functional ribosome pool, and thus, overall protein synthesis (**Fig. 5C bottom right panel**). Additionally, abnormal abundance of specific snoRNAs in the experimental samples might have affected optimal rRNA modification, and that in turn might also have affected the formation of functional ribosomes. Together, it was evident that the endoribonuclease activity of XRN-2 plays a critical role in rRNA metabolism with a direct consequence to the process of translation.

## Discussion

Here, we have functionally characterized the endoribonuclease activity of XRN-2, which demonstrates high fidelity and processivity, while requiring no external energy. Our results suggest that this activity is initiated by the binding of the 5’-phosphorylated end of the substrate miRNA to the PCP (positively charged patch). The PCP comprises nine amino acids, out of that seven are positively charged residues, which are partially conserved. The PCP is likely to be a free energy hotspot, where conserved residues might contribute the maximum, resulting in substantial energy release upon miRNA binding. The released energy is likely to elicit conformational changes in the protein, guiding the substrate miRNA into the catalytic site, which might also get simultaneously organized for metal ion coordination. The energy released from phosphodiester bond cleavage in the catalytic site is likely to drive the movement of the newly formed 5’ end of the RNA towards and into the PCP, replacing and releasing the previously bound 5’-phosphorylated fragment as a cleavage product. Based on the consistent size of the cleavage products (3-nt), it can also be proposed that the PCP and the catalytic site are apart by a distance as much as the length of around three nucleotides. Product formation ceases as soon as the substrate RNA becomes too short that its 3’-end fails to span across the catalytic site, while its 5’-end is presumably bound to the PCP, which in turn facilitates termination of the reaction (**Fig. 3K, reference 12**).

Nuclear 5’-3’ exoribonuclease activity (XRN-2) of eukaryotes is known for over three decades (30, 31), and in the course of time XRN-2 has been shown to be critically important for the processing and turnover of a variety of fundamentally important RNAs, including miRNAs (5-7). But, only recently, its endoribonuclease activity has been identified (12), and here we presented a partial functional characterization of this newly identified enzymatic activity. Interestingly, since its discovery, the eukaryotic exosome complex was known to work in a 3’-5’ exoribonucleolytic mode, until a physiologically important endoribonuclease activity was demonstrated for one of its constituent subunits (Dis3) characterized as a 3’-5’ exoribonuclease (32-34). Therefore, the insights emanating from our studies are highly intriguing, but not completely unprecedented. Additionally, we have presented a novel mechanism of endoribonuclease activity, which requires large-scale conformational changes upon substrate (miRNA) binding. In some ways, it is reminiscent of the formation of the *Kluyveromyces polysporus* Argonaute (KpAGO) catalytic tetrad involving movement of a glutamate residue due to dramatic conformational changes, in otherwise rigid structures, in a guide-bound KpAGO (35).

The residues constituting the endoribonuclease catalytic core of XRN-2 are conserved in higher eukaryotes (except one odd residue in *Drosophila*, **Supplementary figure 6A**), and mutation of the homologous residues in human XRN2 perturbs its endoribonuclease activity *in vitro* (data not shown). Thus, it is possible that this activity is equally important in other eukaryotic organisms, especially vertebrates. But, more interestingly, the PCP residues, which upon binding of substrate elicit conformational changes to organize the catalytic core of the endoribonuclease activity of the worm XRN-2, are only partially conserved (**Supplementary figure 6B**). Here, it is possible that there are some species-specific residues, which critically contribute toward this activity, and they may be targeted for translational purposes. An atomic structure with the substrate RNA-in-place would help us to understand the precise role of the different residues for this enzymatic activity, and may facilitate translational research.

Previous *ex vivo* studies with miRNasome-1 demonstrated its receptiveness for ATP in the environment, where depending on the availability of ATP, it could operate in two alternative modes of catalytic mechanism. Its ATP-independent XRN-2-mediated endoribonuclease activity was shown to be much more efficacious than that of the ATP-dependent exoribonuclease activity. Here, we could establish the physiological significance of the ATP-independent endoribonuclease activity of XRN-2 by demonstrating that it is critical for miRNA metabolism in energy-deprived dauer worms (**Figure 4**), where the levels of some of the substrate mature miRNAs of miRNasome-1 are much diminished compared to that observed in the relevant stages of the continuously growing worms (**Figure 1B**). Accordingly, that justified the higher efficacy of the endoribonuclease activity of miRNasome-1 over its exoribonuclease activity, which gets employed during the dauer stage and responsible for the establishment and maintenance of those concerned miRNAs at such low levels. Although, it was not possible to purify the miRNasome-1 from dauer worms due to limitation in the amount of extracted protein, but using co-immunoprecipitation experiments, we could find that the different components of miRNasome-1 indeed interact with each other in the dauer worms, albeit, miRNasome-1.4 showed very weak interaction (data not shown). It suggested that miRNasome-1 might exist in the dauer worms as an intact complex, which in turn may allow the organism to quickly switch to the energy dependent exoribonucleolytic mode of miRNA turnover mechanism upon arrival of favourable conditions. Notably, the product of the exoribonucleolytic turnover mechanism, monoribonucleotides, is likely to get reused in different synthetic pathways facilitating growth and development.

Perturbation of the newly identified endoribonuclease activity of XRN-2 proved to be fatal for the sturdy quiescent dauer worms (**Fig. 4A**). In those experimental samples, we observed downregulation of many dauer specific genes, especially transcription factors, metabolic enzymes and alpha-crystallin family of heat shock proteins that are critical for dauer maintenance (data not shown). It was quite apparent that a number of critical pathways did fail in those experimental dauers, and culminated in their fall. Since, the dauer quiescence is analogous to that of the quiescent stage of the adult stem cells in higher eukaryotes and mammals (**15, 16**), and to the infective stage of the parasitic nematodes (**13**), further insights on the entire spectrum of substrate miRNAs of this activity, as well as the effects on the target-mRNAs and downstream events upon perturbation of this activity might facilitate a better understanding of the biology of quiescent adult stem cells, and the parasitic nematodes in their infective stages.

Genetic study-driven inferences, not involving specific mutants of the exoribonuclease activity of XRN-2, can raise questions on the correctness for attribution of those many roles to XRN-2’s ‘default’ exoribonuclease activity, when it is possible that some or many of those different roles in different pathways are actually played fully or partially by the newly identified endoribonuclease activity. Indeed, we could demonstrate that endoribonuclease activity of XRN-2 is important for the biogenesis of mature 28s and 5.8s rRNAs by removal of precursor sequences (internal transcribed spacers) of the pre-rRNA. But, our work does not exclude the role of the exoribonuclease activity of XRN-2 on rRNA metabolism, and it is possible that orchestration between both the activities do take place. More work needs to be done to understand the pre-rRNA sequence and structure requirements, as well as the role of any other partner proteins, which might determine and regulate specific endoribonucleolytic cleavages on precursor molecules. Notably, rRNAs in the dauer worms undergoing perturbation of the endoribonuclease activity of XRN-2 remained unaffected (data not shown), and it is possible that the selected early time point was too early to record an effect on the stable rRNAs. Importantly, this data further strengthened the message that the deregulation of miRNAs and their targets in dauers were specific consequences of the perturbation of the endoribonuclease activity of XRN-2, and not due to any indirect effect. Additionally, continuously growing *xrn-2*(*PHX25*) worms mutant for the endoribonuclease activity of XRN-2 exhibited a conditional mortal germline (mrt) phenotype (**Supplementary figure 4A)**, when they were grown at the non-permissive temperature (26°C). It is known that worms with mutations in protein(s) involved in the pathways governing nuclear RNAi (eg. NRDE-2) and trans-generational inheritance of epigenetic memory (eg. HRDE-1) demonstrate similar mrt phenotype (36-38). So it is possible that perturbation of the endoribonuclease activity of XRN-2 directly or indirectly affected those pathways, and thus, in turn led to the sterility of the organisms.

To conclude, further identification of other endogenous RNA substrates, followed by detailed understanding of the mechanistic roles of this endoribonuclease activity on those transcripts through both *in vivo* and *in vitro* studies would harness critical new insights. Additionally, understanding of the regulation of those multiple roles played by this activity under different physiological conditions would bring about revelations that would certainly be important to appreciate the biogenesis and degradation of different species of RNA molecules. Finally, if this new enzymatic activity is conserved across eukaryotes, it can essentially act as a platform for a new wave of fundamental research and paradigm shift in the broader RNA research field.

## Author Contributions

SC designed the research and experiments. MS, AS, ARW performed the experiments and together all the authors analyzed the results. SC wrote the manuscript.

## Acknowledgments

We thank Mohini Singh for technical help. This work has been supported by the Wellcome Trust-DBT India Alliance through a fellowship to SC. MS and AS received fellowships from UGC and CSIR (Govt. of India), respectively. ARW received support from SERB/ DST (Govt. of India).

