## Supplementary Information for "Endoribonuclease activity of XRN-2 is critical for RNA metabolism and survival of *Caenorhabditis elegans*"

**This file includes:**
Materials and Methods
Supplementary figures 1 to 6 with their captions
Supplementary Table 1

### Materials and Methods

**Worm strains and RNAi.** The wild type strain *C. elegans* var. Bristol strain *N2* has been used. The other strain was *xrn-2*(*xe31*) (a temperature sensitive *xrn-2* mutant strain harboring a point mutation in the *xrn-2* gene, reference 39). All phenotypic readouts were an average from three biological replicates ± SEM, where n = 200, if not otherwise indicated. RNAi was performed as described before (12).

**Generation of a worm strain mutant for the endoribonuclease activity of XRN-2.** The mutant strain *xrn-2*(*PHX25)* of the *xrn-2*(*syb25*) allele, which has base substitutions in the *xrn-2* coding sequence leading to D86-A and D181-A, was generated by SunyBiotech (PRC) using the CRISPR/ Cas9 technology. This strain [*xrn-2(PHX25*)] houses the two aforementioned mutated residues out of the five active site residues of the endoribonucleolytic activity of XRN-2. The strain was homozygoused, and verified by DNA sequencing.

**RNA isolation, Northern blotting, RT-qPCR, TaqMan analysis.** Total RNA was isolated from staged L4 worms, using Trizol (Ambion) method as described before (40). Northern blotting of endogenous RNA was done as described (41). 5'-labeled DNA oligos were used as probes, where OptiKinase™ (Phosphatase minus mutant of T4 polynucleotide kinase [PNK, Affymetrix]) and γ-$^{32}$P-ATP were used to perform the 5'-end labeling. The hybridizations for the members of the *let-7* family of miRNAs were carried out at an elevated temperature of 40°C in order to minimize the binding of the probe to unintended *let-7* sisters (5). For RT-qPCR, 2 µg of total RNA of the respective samples, was reverse transcribed using SuperScript™ III Reverse Transcriptase (Invitrogen) in 20 µl reaction containing 5 mM DTT, 0.5 mM dNTPs, 2.5 µM oligo

$(dT)_{20}$ and 1 μl of enzyme, in accord with the manufacturer's protocol. 1.5 μl of each of the reverse-transcription reactions were amplified with PowerUp™ SYBR Green Master Mix (Applied Biosystems) in a total volume of 10 μl using specific forward and reverse primers at a concentration of 0.5 μM each, and analyzed in a ViiA™ 7 Real-Time PCR System using ΔΔCt method. The primers used are described below. Values were normalized against the value of act-1 expression. The experimental values, representing means (±SEM) from three independent biological replicates, were compared to the expression levels of the corresponding *N2/ xrn-2(xe31)/ xrn-2(PHX25)* (Empty Vector/ Control) samples, which were set as 1. TaqMan MicroRNA Assays (Applied Biosystems) to quantify individual miRNAs were performed as per the supplier's protocol and as described (42). In brief, 10 ng of total RNA of the respective samples, was reverse transcribed in 7.5 μl reaction volume and then 0.67 μl of each of the reverse-transcription reactions was amplified in a total volume of 10 μl, and analyzed in a ViiA™ 7 Real-Time PCR System using ΔΔCt method. Values were normalized against the value of *sn2841* and *U18* expression in the respective samples.

**Cloning and expression of recombinant XRN-2.** cDNA was generated from *C. elegans* total RNA using SuperScript™ III Reverse Transcriptase (Invitrogen) and oligo $(dT)_{20}$. Then *xrn-2* cDNA was PCR amplified from the cDNA using gene specific primers and Q5 High-Fidelity DNA Polymerase (NEB), and cloned in a TOPO TA vector (Invitrogen). The sequence confirmed correct ORF was subcloned in pFastBac™ HT B (Invitrogen), and further used to generate the recombinant bacmid DNA as per the manufacturer's protocol (Bac-to-Bac Baculovirus Expression System, Invitrogen), towards expression of the recombinant N-terminal Strep tag containing wild type, as well as the different mutant XRN-2 proteins. They were purified using Strep-Tactin$^{R}$XT Superflow$^{R}$ high capacity resin (IBA, Germany). Of note, high salt (0.25-0.5M) containing wash and elution buffers were used for the removal of nonspecific host proteins and for the complete elution of the recombinant protein, respectively. All the above proteins were subjected to dialysis to remove the excess salt before performing any functional assay.

**Generation of recombinant truncated proteins and proteins with substituted amino acids.** Truncated versions of the *C. elegans xrn-2* [Ce-XRN-2 N-terminal fragment (1-474 aa), Ce-XRN-2 Mid fragment (475-706 aa), Ce-XRN-2 C-terminal fragment (707-975 aa)] were amplified using specific primers, cloned in pGEX4T-1 expression vector (GE Healthcare), and sequence confirmed. Point mutations were introduced in the full length as well as the truncated XRN-2 N-terminal fragment (1-474 aa) through site directed mutagenesis, following the protocol detailed in the QuikChange site directed mutagenesis kit (Agilent). In some cases, DNA fragments of *xrn-2* gene with necessary mutations (for potential endoribonuclease active site residue clusters and positively charged patch residues) were obtained from GenScript (USA), and then cloned in appropriate vectors. All the clones were sequence confirmed.

**Preparation of RNA substrates.** Mature *let-7* and mature *mir-84* RNA were prepared as per the methods described before (8, 12, 43). In brief, a chimeric RNA comprising of a hammerhead ribozyme in its 5'-end followed by the mature *let-7/ mir-84* sequence was transcribed from DNA cassettes using a MEGAshortscript$^{TM}$ T7 transcription kit (Ambion) in presence of $\alpha$-$^{32}$P-UTP, as per the supplier's protocol. The DNA cassettes were prepared by annealing of appropriate forward and reverse primers (please see the oligo sequence section) followed by fill-in reactions using Klenow (NEB). Double stranded DNAs of appropriate length were gel purified and directly used as template for *in vitro* transcription. The gel purified products were also PCR amplified, cloned and sequence confirmed. While transcription reaction is ongoing, self-processing of the ribozyme-containing transcripts occur simultaneously. The resulting mature *let-7/ mir-84* containing 5' hydroxyl groups were size purified by 7 M urea/ 8-10% PAGE. After recovery RNAs were 5' phosphorylated by T4 PNK (NEB) and ATP.

5' labeling of mature miRNAs (Microsynth AG, Switzerland) were done using T4 PNK (NEB) and $\gamma$-$^{32}$P ATP. Before use the radiolabeled synthetic pre-*let-7* RNA was subjected to refolding as described before (8, 12, 43). 3' labeling and blocking of synthetic mature miRNAs were done using T4 RNA ligase (Ambion) and [5'-$^{32}$P] pCp, according to manufacturer's instructions. All RNAs were size purified using 7 M urea/ 8-10% PAGE.

**Preparation of worm lysate.** Staged L4 worms were grown on plates and harvested with M9, and washed thrice with the same buffer to eliminate any bacteria. The worm pellet was then resuspended in extraction buffer (25 mM HEPES [pH 7.4], 2.5 mM DTT, 2.5 mM $MgCl_2$, 0.1% Triton-X 100, 100 mM KCl, 10% Glycerol, 1X SIGMAFAST$^{TM}$ Protease Inhibitor Cocktail) and ground in liquid $N_2$. The clear supernatant was collected after spinning the thawed sample at 16,000 g for 30 min, which was further sequentially passed through Miracloth (Calbiochem), empty Poly-Prep Chromatography Column (Bio-Rad), and 0.4 micron filter unit (Millipore), and designated as 'total lysate/ worm lysate' or simply 'lysate'. Of note, cleared worm lysate was used in all the *in vitro* assays involving lysate.

***in vitro* turnover assay.** Labeled RNAs (mature *let-7*/ mature *mir-84*) approximately 100 fmol, if not otherwise indicated, were incubated with cleared worm lysate (1-5 μg) or a given recombinant protein (10-100 ng) in 1X assay buffer (AB; 25 mM HEPES [pH 7.4], 2.5 mM DTT, 5 mM $MgCl_2$, 100 mM KCl, 2.5 mM ATP) in a volume of 10 μl at 25°C for 15 min. The reactions were terminated by addition of 1X volume formamide gel loading buffer (95% Formamide, 0.2% SDS, 1 mM EDTA, 0.04% Xylene Cyanol, 0.04% Bromophenol Blue) followed by heating at 65°C for 5 min. Equal volumes of the samples were then subjected to 7 M urea/8-16% PAGE followed by gel drying and autoradiography or phosphor-imaging.

The RNA size ladders for *mir-84* and *let-7* were generated as per the Ambion's partial alkaline hydrolysis protocol, where a 5'-radiolabeled RNA (*mir-84, let-7*) was subjected to incomplete hydrolysis by alkali.

**Assay determining the mode of catalysis employed by the endoribonuclease activity of XRN-2 (Metal ion-mediated vs Acid-base catalysis).** Towards this goal, 5 nM (110 ng) purified recombinant wild type XRN-2 was incubated with 15 nM (300 fmol) of 5'-radiolabeled *let-7* as a substrate in the AB in a volume of 20 μl at 25°C for 15 min, in parallel replicates. These reactions were terminated, phenol-chloroform extracted, and alcohol precipitated. Then one replicate was subjected to 5'-end labeling using [γ32P] ATP (3500 Ci/mmol) and T4 polynucleotide kinase (NEB) in a total volume of 15 μl at 37°C for 45 min. Equal volumes of the samples were then subjected to 7 M urea/12% PAGE followed by gel drying and autoradiography or phosphor-imaging. Any increase in the signal for XRN-2- product would be indicative of the endoribonuclease mechanism involving acid-base catalysis, which results in the formation of 5'-OH products, and thus allows subsequent 5'- phosphorylation. Contrastingly, no increase in signal for XRN-2-products would be indicative of a metal ion-mediated mechanism that produces products with 5'-phosphates, which would not be further phosphorylated. Similar reactions and subsequent analyses were performed with a 5'-radiolabeled *let-7* and 1 ng of Ribonuclease A (RNase A), which served as a positive control for acid-base mode of catalysis.

**Antibodies and Western blotting.** Rat α-XRN-2 serum was generously provided by Helge Grosshans (44). For western blotting, α-XRN-2 was used at 1: 2000 dilutions, followed by the use of α-Rabbit HRP-conjugated secondary antibody (GE Healthcare) at a dilution of 1: 10,000. Detection was performed with Amersham™ ECL™ Prime Western Blotting Detection Reagent, and Amersham Hyperfilm™ ECL or ImageQuant LAS 4000 Chemiluminescence Imager (GE Healthcare). Band sizes (relative molecular mass) were determined using GS-900™ software (Bio-Rad).

**Endogenous ATP measurement.** ATP was measured by ATPlite Luminescence ATP Detection Assay System (PerkinElmer) according to the manufacturer's protocol. Briefly, equal pack-volume of worms were harvested at specific stages and mixed with equal volume of the provided lysis solution, and then ground in liquid nitrogen to prepare the total lysates. For measurement, 7.5 μl of lysate from each sample was taken in a tube followed by addition of 7.5 μl of the reconstituted substrate from the kit, and shaken for 5 minutes. Luminescence was measured with TD-20/20n Luminometer (Turner Designs) only after 10 minutes of dark adaptation. The readings were converted to counts per minute and normalized against per microgram of protein.

**XRN-2 inactivation in dauer worms.** Temperature sensitive *xrn-2* mutant worms (*xe31*, reference 39) were pushed into dauer stage through starvation and over-crowding. Then randomly ten worms were picked from each plate for microscopic confirmation for the presence of mouth-plug and atrophied pharynx, which are distinctive features of dauer worms. Plates from where at least seven out of ten worms tested positive for the aforementioned dauer phenotypes were selected, and further maintained for four more weeks. To perturb XRN-2, half of those starved dauer worm plates were shifted to the non-permissive temperature of 26ºC (experimental dauer worms), whereas the other half was maintained at the permissive temperature of 20ºC (control dauer worms). Dead worms were counted at 12 hr intervals, and removed with a platinum wire. The experiment was repeated three times. Survival analysis (n = 2000) was performed using Kaplan-Meier estimate, and the survival curve was plotted using GraphPad Prism. After 48 hrs of dwelling under the two different conditions (permissive vs non-permissive temperature), the control and experimental dauer worms were harvested and subjected to 1% SDS treatment for 30 minutes to eliminate any non-dauer worm, and further subjected to several molecular analyses (status of XRN-2 protein, miRNA levels, validated target mRNA levels). After SDS treatment, small aliquots were plated to confirm the absence of any dead 'non-dauer' worm.

**Inactivation of the endoribonuclease activity of XRN-2.** The generated mutant strain [*xrn-2*(*PHX25*)] having two mutated residues out of the five active site residues of the endoribonuclease activity of XRN-2 was expected to have diminished activity. It was found to be a temperature sensitive mutant, as it grew normally in the permissive temperature (20ºC), but showed a number of phenotypic defects at the non-permissive temperature (26ºC). Dauers of this strain were also generated as described above. They survived at the permissive temperature (20ºC) for more than 50 days, but succumbed within 10 days, when maintained at the non-permissive temperature of 26ºC.

**Recording of the phenotypic defects in the temperature sensitive mutant strain *xrn-2*(*PHX25*) for the endoribonuclease activity of XRN-2.** The experiment was started with two sets of synchronized *xrn-2*(*PHX25*) L1 worms (each set comprising of ten plates with a single worm on each plate), one maintained at the permissive 20°C and the other one at the non-permissive 26°C. Gravid embryo-laying worms were moved to fresh plates after every 12 hours. The total number of embryos laid by each worm was counted, and subsequently the numbers of hatched L1 worms were also recorded. One such L1 worm was picked from each of the plates from the two sets, and further maintained in fresh plates under the same conditions. This was repeated for every new generation until the worms at the non-permissive temperature arrested or died . For recording of phenotypes other than brood size and number of embryos hatched, experiments were started with 500 synchronized L1 worms (P0). When they became gravid, they were bleached to isolate the embryos, from which synchronized L1 worms were generated. 500 of them (F1) were used to continue the experiment. This was continued until F9. The phenotypes were recorded in every generation. The entire experiment was also performed with *N2* worms as an additional control, but they did not show any aberrant phenotype. All the aforementioned experiments were repeated three times.

**Thermal Shift Assay.** In a 20 µl reaction, 450 nM recombinant XRN-2 protein was mixed with Protein Thermal Shift™ Dye and Buffer (Applied Biosystems, USA) to a final concentration of 1X. 5'-phosphorylated synthetic *let-7* miRNA (Microsynth AG,

Switzerland) was added to the reaction to a final concentration of 45 μM for testing the effect of substrate binding. The reactions were set in MicroAmp® optical microplate at 4°C and subjected to a melt curve program from 25°C – 99°C, as per the manufacturer's instructions using a Viia™ 7 real-time PCR system. Raw data was exported to Protein Thermal Shift™ Software, which generated the melting temperature (*Tm*) values from the melt curves by employing the Boltzmann-derived *Tm* method, and it also determined the Δ*Tm* values (shift in *Tm* between the only protein and protein + miRNA melt curves). To plot the graph in Microsoft excel, the resulting sigmoidal part of the curve was fit to Boltzmann Equation using non-linear fitting, in order to show the *Tm* as the midpoint of the unfolding transition.

**Polysome Analysis.** Worms grown on OP50 seeded NGM plates were harvested with M9, and washed thrice with cold M9, and once with cold autoclaved glass distilled water. Worm pellets were then resuspended in 300 μl of cold lysis buffer [20 mM Tris-HCl (pH 8.5), 140 mM KCl, 1.5 mM MgCl2, 0.5% Nonidet P40, 2% PTE (polyoxyethylene-10-tridecylether), 1% DOC (sodium deoxycholate monohydrate), 1 mM DTT, 1X SIGMAFASTTM Protease Inhibitor Cocktail] and were crushed to a fine powder using pre-cooled mortar and pestle. After the powder thawed, lysates were collected and cleared by centrifugation (10 min. at 10,000 g, 4°C). Absorbance of the resulting cleared lysate at 260 nm were measured and equivalent amounts of control and experimental samples were loaded onto 12 ml 15-60% sucrose gradient in gradient buffer [20 mM Tris-HCl (pH 8.5), 140 mM KCl, 1.5 mM MgCl2 and 1 mM DTT], and centrifuged in a Beckman SW41Ti rotor at 39,000 rpm at 4°C for 3 hours.

Gradients were then fractionated from top to bottom into 20 fractions of equal volume using a BIOCOMP density gradient fractionator with continuous monitoring of absorbance at 260 nm to detect free-fraction, ribosomal subunits, monosomes and polysomes. RNA from each fraction was extracted using TRIzol (Invitrogen), according to the manufacturer's instructions.

**CD Spectroscopy.** To confirm proper protein folding of the wild type XRN-2 and other mutants CD spectra (wavelength 190–260 nm) were recorded with a JASCO 815 CD Spectrometer at 5˚C in a 0.1-cm cuvette. Proteins were diluted in buffer containing 1 X PBS and 2 mM DTT to a final protein concentration of 2 μM in a total volume of 300 ul. Five scans were taken with a speed of 200 nm/min.

**Structure homology modelling and analysis.** Structure homology was computed using SWISS-MODEL (Swiss Institute of Bioinformatics, Biozentrum, University of Basel, Switzerland) homology server (45). Model for full length C. elegans XRN-2 was built using 5fiR.pdb (18) as template by employing SWISS-MODEL. All figures and structures were plotted and visualised, respectively, using PyMOL (Schrodinger LLC; Educational-use-only), UCSF Chimera (46) and Chimera X (47). Structural comparisons between different PDB files were performed by using the MatchMaker extension of Chimera X. The MatchMaker extension of Chimera constructs pairwise sequence alignments and uses them to superimpose the structures. Following which, a full set of residue equivalences between the structures were obtained using Match -> Align extension of Chimera X.

**Oligos (5’-3’).**

<u>Northern</u>:

*let-7*(WT): AAC TAT ACA ACC TAC TAC CTC A

*mir-84*: TCT ACA ATA TTA CAT ACT ACC TCA

*mir-48*: TCG CAT CTA CTG AGC CTA CCT CA

*mir-241*: TCA TTT CTC GCA CCT ACC TCA

*mir-77*: TGG ACA GCT ATG GCC TGA TGA A

*mir-237*: AGC TGT TCG AGA ATT CTC AGG GA

*lin-4*: TCA CAC TTG AGG TCT CAG GGA

*mir-90*: AGG GGC ATT CAA ACA ACA TAT CA

*mir-124*: TGG CAT TCA CCG CGT GCC TTA

*mir-75*: TGA AGC CGG TTG GTA GCT TTA A

*mir-1*: TAC ATA CTT CTT TAC ATT CCA

*mir-234*: AAG GGT ATT CTC GAG CAA TAA

*mir-79*: AGC TTT GGT AAC CTA GCT TTA T

5.8S rRNA : CAA CCC TGA ACC AGA CGT ACC AAC TGG AGG CCC AGT TGG T

26S rRNA : CGG TAC TTG TTC GCT ATC GCA ATC GAG TCG ATA TTT AGC

18S rRNA : GGT TCA CCT ACA GCT ACC TTG TTA CGA CTT TTA CCC G

$tRNA^{Gly}$: GCT TGG AAG GCA TCC ATG CTG ACC ATT

<u>TaqMan microRNA Assays (Applied Biosystems) were used for the following miRNAs and control RNA:</u>

*let-7*(WT), *mir-84, mir-48, mir-241, mir-77*, *mir-237, lin-4, mir-90, mir-124, mir-75, mir-1, mir-234, mir-79, mir-57, mir-71, mir-230, mir-54, mir-788, mir-795, mir-83, sn2841* qPCR:

Primary *mir-241*

Forward: CAT CCT TCC GCT TGT TGT TT

Reverse: ACC CAT TCA ACC AAA ATC CA

Primary *mir-84*

Forward: TGT CTG GTT TCG GCG GAT AG

Reverse: CCA CAG GCA GAC GTA TGA TG

Primary *mir-90*

Forward: GTC GTC CTT CTT TTC CAC CAG

Reverse: GTC GGT TGA TAT CGT CGT TG

*daf-12* mRNA

Forward: GAT CCT CCG ATG AAC GAA AA

Reverse: CTC TTC GGC TTC ACC AGA AC

*hbl-1* mRNA

Forward: ATG GTG CAA TCC GAT AGT CC

Reverse: TGG TAA TTT GAA GAT GTG CCC TC

*act-1* mRNA

Forward: CAC GGT ATC GTC ACC AAC TG

Reverse: GTA CGT CCG GAA GCG TAG AG

Cloning:

*xrn-2* cDNA (for Baculovirus expression)

Forward Primer: CAA GAA TTC TAA TGG GAG TTC CCG CAT TCT TCA GAT G

Reverse primer: GCG AAG CTT TTA TCT CCA TGA TGA ATT TCC GTG ATA G

XRN-2 N-terminal fragment (1-474 aa)

Forward Primer: GGA GAA TTC ATG GGA GTT CCC GCA TTC TTC AG

Reverse Primer: GAA GCG GCC GCT TAT TCC GAG GCC ATC TGC C

XRN-2 Mid fragment (475-706 aa)

Forward Primer: GAA GAATTC GCC AGG CAG ACG GCC ATG

Reverse Primer: GAA GCGGCCGC TTA TCC ACG TGT ATT TCG TTG CTT TTC

XRN-2 C-terminal fragment (707-976 aa)

Forward Primer: GGA GAATTC CCG AAT CGA ATT TTC ATT GGA CG

Reverse Primer: GAA GCGGCCGC TTA TCT CCA TGA TGA ATT TCC GTG

<u>Site Directed Mutagenesis (SDM):</u>

XRN-2 D37A

Forward Primer: GGG CTG TGT ACA AGC TAC GGG GAC CCG

Reverse Primer: CGG GTC CCC GTA GCT TGT ACA CAG CCC

XRN-2 D86A

Forward Primer: TGA CAA TCG AAT AAA TGC GAG CGA TGT ACT CGA AAA TGA GC

Reverse Primer: GCT CAT TTT CGA GTA CAT CGC TCG CAT TTA TTC GAT TGT CA

XRN-2 D181A

Forward Primer: GCG TCA TTA GTG ACC CGA GCA TGA ATA TAA TAT CGA AGA G

Reverse Primer: CTC TTC GAT ATT ATA TTC ATG CTC GGG TCA CTA ATG ACG C

XRN-2 D186A

Forward Primer: TAG CCC ATG ACG CGG CAT TAG TGA CCC GAT C

Reverse Primer: GAT CGG GTC ACT AAT GCC GCG TCA TGG GCT A

XRN-2 E23A

Forward Primer: CGG CCT GGG CAT CTG CAT TGG CAT TCA CC

Reverse Primer: GGT GAA TGC CAA TGC AGA TGC CCA GGC CG

Preparation of templates for *in vitro* transcription:

Mature *mir-84* cassette:

Forward primer (T7 Promoter, HH Ribozyme, First 12 *mir-84* nt.)
G TAA TAC GAC TCA CTA TAG GG AGA CAT ACT ACC TCA CTG ATG AGT CCG TGA GGA CGA AAC GGT ACC CGG TAC CGT CTG AGG TAG TAT G

Reverse primer (Mature *mir-84* complementary sequence, 12 nt. complementary region to HH Ribozyme)
TCT ACA ATA TTA CAT ACT ACC TCA GAC GGT ACC GGG

Mature *let-7* cassette:

Forward primer (T7 Promoter, HH Ribozyme, First 12 *let-7* nt.)
G TAA TAC GAC TCA CTA TAG GGAGA CTA CTA CCT CAC TGA TGA GTC CGT GAG GAC GAA ACG GTA CCC GGT ACC GTC TGA GGT AGT AGG

Reverse primer (Mature *let-7* complementary sequence, 12 nt. complementary region to HH Ribozyme)
AAC TAT ACA ACC TAC TAC CTC A GAC GGT ACC GGG

Mature *mir-237* cassette:

Forward primer (T7 Promoter, HH Ribozyme, First 13 *miR-237* nt.)
G TAA TAC GAC TCA CTA TAG GGG AGA CGA GAA TTC TCA GGG ACT GAT AGT CCG TGA GGA CGA AAC GGT ACC CGG TAC CGT CTC CCT GAG AAT TCT CGA ACA GCT

Reverse primer (Mature *mir-237*, 13 nt. Complementary region to HH Ribozyme)
AGC TGT TCG AGA ATT CTC AGG GA GAC GGT ACC GGG

### Synthetic RNAs (5'-3').

*let-7*: UGA GGU AGU AGG UUG UAU AGU U

*mir-84*: UGA GGU AGU AUG UAA UAU UGU AGA

*lin-4*: UCC CUG AGA CCU CAA GUG UGA

### Supplementary figures

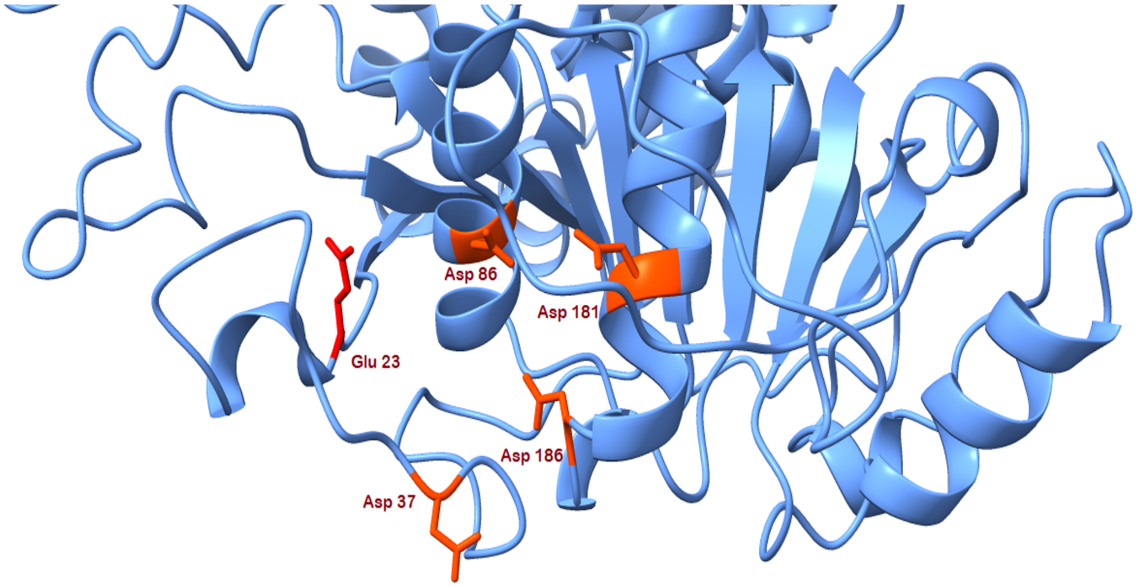


**Supplementary figure 1. Glu23 (red) of XRN-2 is positioned in close proximity to the cluster II residues as depicted in the image (orange).** Glu23 and Asp86 are 11.5 Å apart.

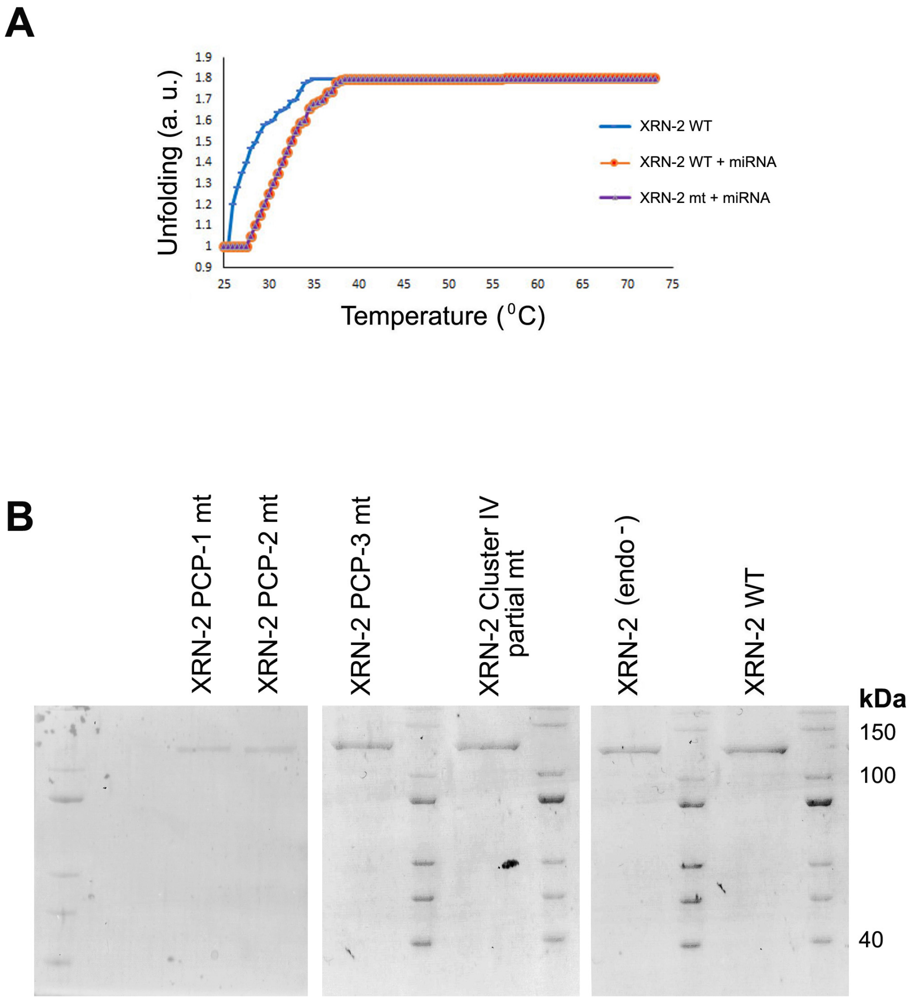


**Supplementary figure 2. (A)** In the presence of substrate miRNA, recombinant wild type (WT) and mutant (mt, all active site residues mutated) XRN-2 melt at a higher temperature to unfold for dye binding, compared to that of the wild type protein alone. Thermal shift assay performed with the indicated proteins in the absence/ presence of miRNA.

**(B)** Wild type and different mutant proteins, as indicated, used for the above experiment and elsewhere in this study.

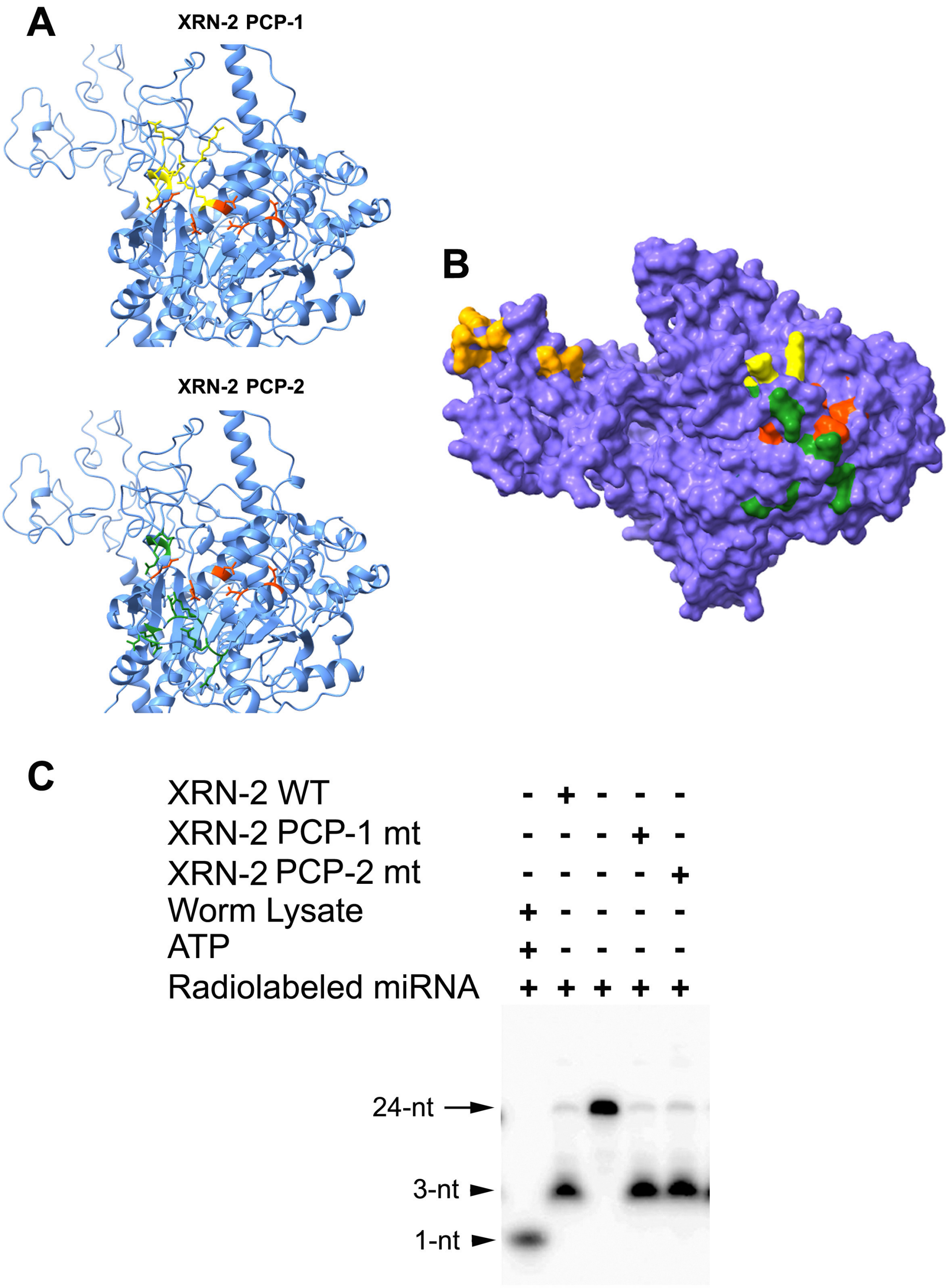


**Supplementary figure 3. Surface-localized positively charged residues near the endoribonuclease catalytic core do not form the binding site for the miRNA**

**substrate. (A)** Zoomed-in views representing the two sets of putative RNA-interacting residues forming two distinct positively charged patches (PCP). PCP-1 (highlighted in yellow) comprises R25, R27, Q29, R33, R87, K298 and PCP-2 (highlighted in green) comprises R25, R27, Q40, Q46, R93, R95, R96, R304.

**(B)** Surface representation of the XRN-2 full length protein, depicted via a space-filling model, indicating the active site residues (red), and the two sets of potential RNA-interacting amino acid residues, PCP-1 (yellow) and PCP-2 (green). A third set of surface-localized residues forming a potential positively charged patch for binding RNA are highlighted in brown (PCP-3).

**(C)** Purified wild type and mutant proteins (XRN-2 PCP-1 mt, XRN-2 PCP-2 mt; 10 nM of each protein) were incubated with radiolabeled substrate miRNA at 25ºC for 15 minutes and resolved on 10% urea-PAGE. The mutations do not affect the endoribonuclease activity of XRN-2.

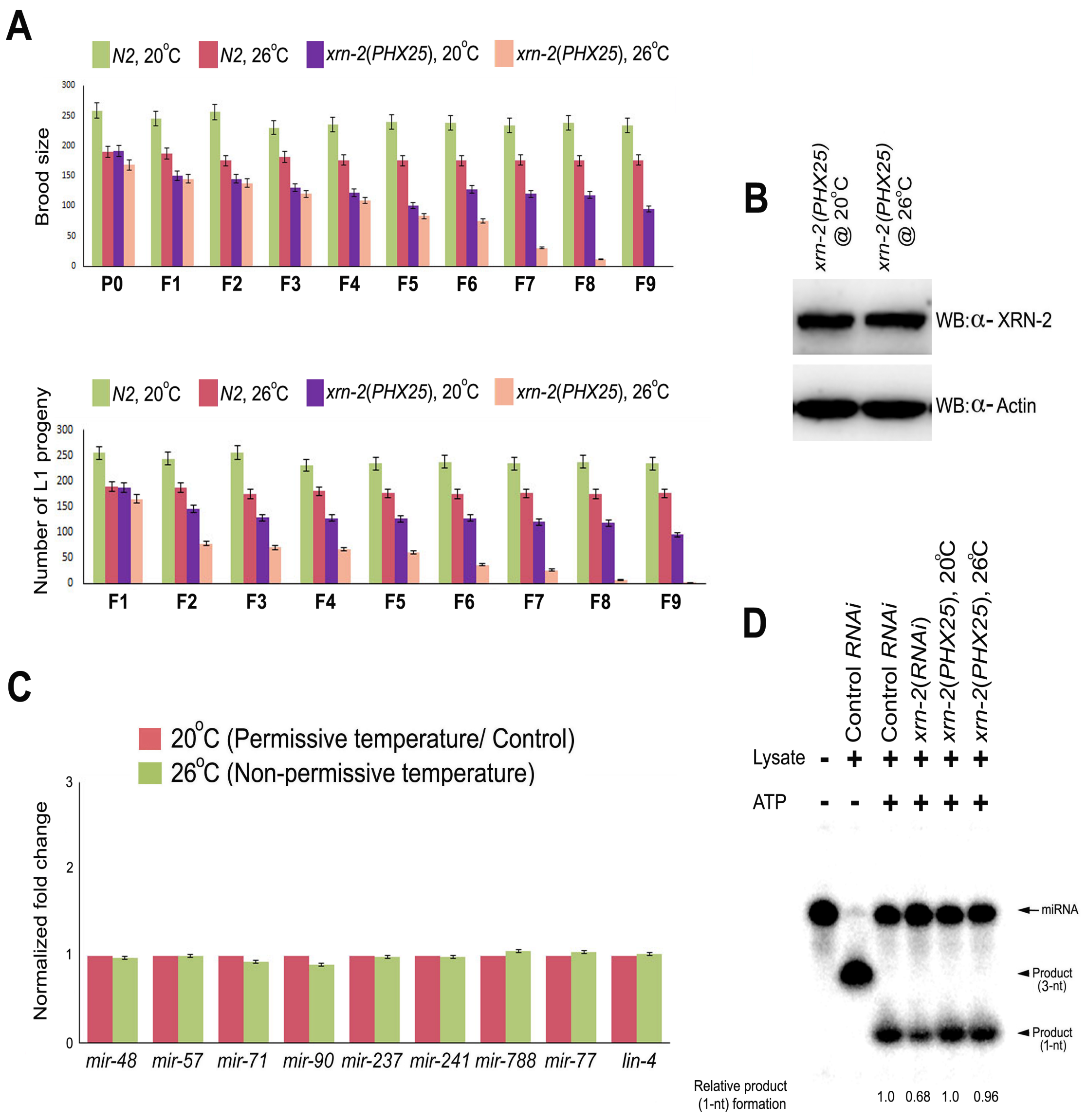


**Supplementary figure 4. Endoribonuclease activity of XRN-2 is important for progeny formation in continuously growing worms, but not for the maintenance of mature miRNA levels.**

**(A)** Mutation in the endoribonuclease active site of XRN-2 results in reduced brood size (**top panel**), and reduction in the number of hatched L1 worms (**bottom panel**) at the non-permissive temperature of 26°C.

**(B)** Western blotting reveals similar XRN-2 levels in both control and experimental worms at the L4 stage of the fourth generation.

**(C)** Mature miRNA levels remain unaffected in the experimental worms at the L4 stage of the fourth generation compared to that of the control worms at the same generation and stage.

**(D)** Lysates from *xrn-2*(*PHX25*) worms growing at the permissive and non-permissive temperatures show equivalent XRN-2-dependent and ATP driven exoribonucleolytic product (monoribonucleotide) forming activity in *in vitro* miRNA turnover assay.

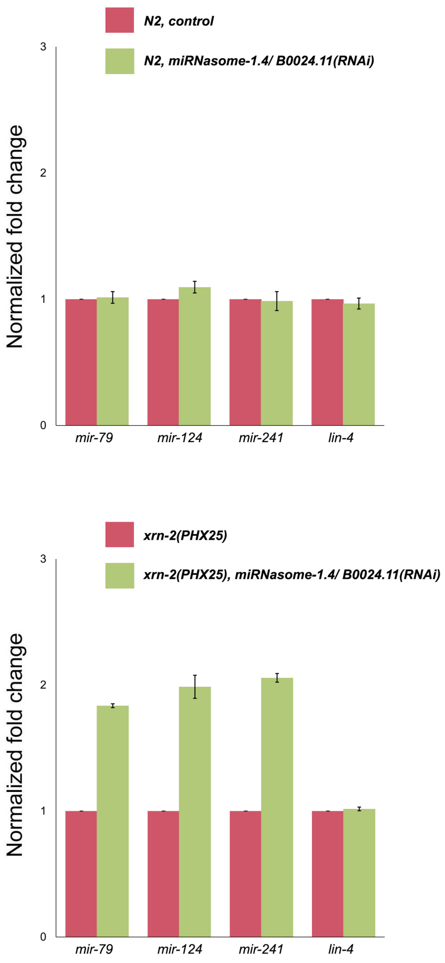


**Supplementary figure 5. Endoribonuclease activity of XRN-2 confers dynamism to the mechanism of miRNA turnover in worms. Top panel.** RNAi depletion of miRNasome-1.4/ B0024.11 has no effect on miRNasome-1 substrate mature miRNA levels in *N2* worms. **Bottom panel.** Depletion of miRNasome-1.4/ B0024.11 results in the accumulation of miRNasome-1 substrate mature miRNAs in *xrn-2*(*PHX25*) worms, where the endoribonuclease activity of XRN-2 is compromised.

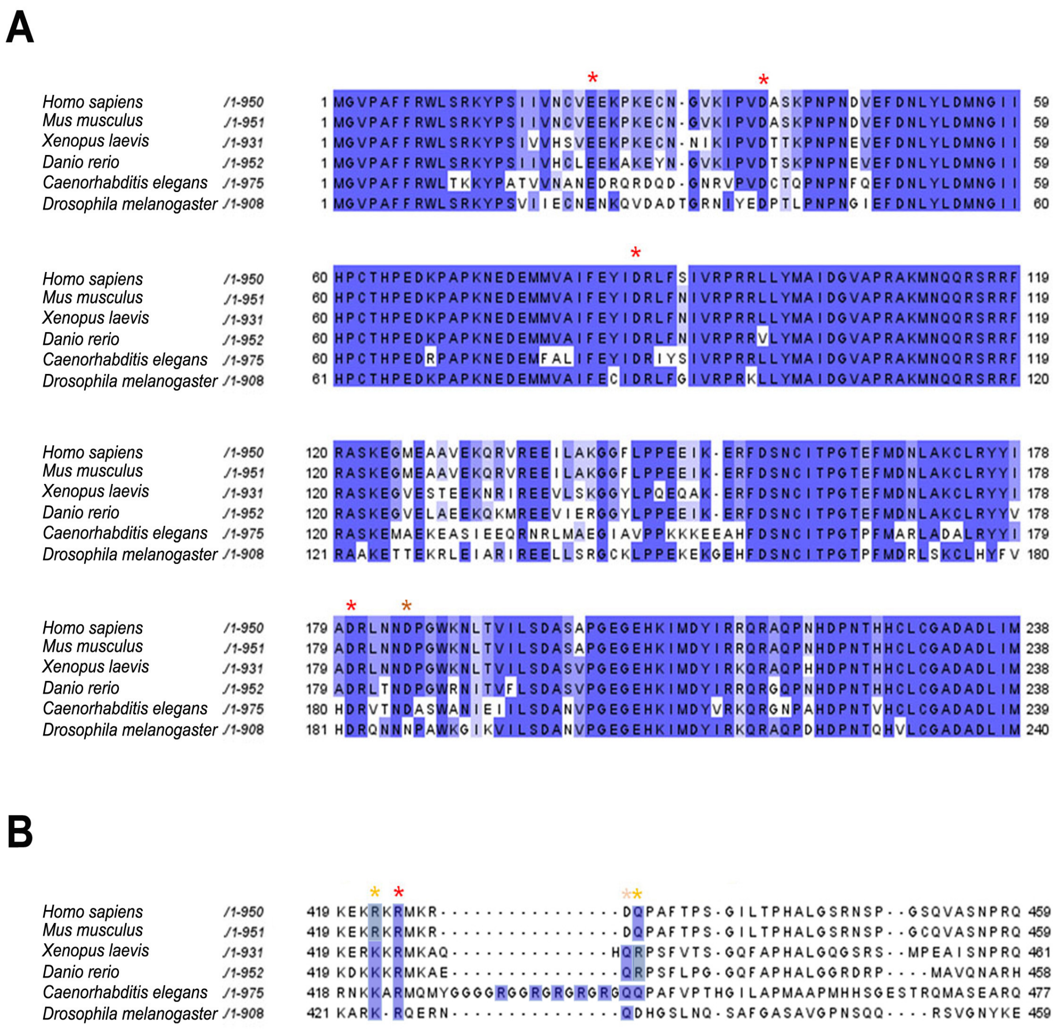


**Supplementary figure 6. (A)** Sequence alignment of XRN-2 protein from different species, as indicated, demonstrates conservation of active site residues in higher eukaryotes, especially vertebrates. Conserved amino acid residues are highlighted in dark blue and similar residues in light blue.

**(B)** Only a partial conservation is observed for the positively charged patch (PCP) residues.

| Phenotypes | Generations | | | | | | | | | |
|---|---|---|---|---|---|---|---|---|---|---|
| | P0 | F1 | F2 | F3 | F4 | F5 | F6 | F7 | F8 | F9 |
| **Shortened and deformed pharynx*** | | | 20% ± 1% | 25% ± 1.5% | 42% ± 2% | 55% ± 2% | 60% ± 2% | 65% ± 2% | 65% ± 1% | 80% ± 1% ** |
| **Defect in vulval development** | | | 5% ± 1% | 20% ± 2% | 40% ± 2.5% | 70% ± 2% | 70% ± 1% | 71% ± 2% | 70% ± 2% | |
| **Reduced brood size** | | 11.5% ± 8.684% | 32.2% ± 7.5% | 45% ± 11.2% | 57% ± 8.75% | 69.5% ±10.51 % | 73% ± 10.12% | 88% ±7.89 % | 90% ± 5% | |
| **Shorter body length*** | | | | 25% ± 5% | 40% ± 2.5% | 41% ± 2.5% | 40% ± 2% | 40% ± 2% | 47% ± 1% | 75% ± 1% ** |
| **Bent posterior** | | 92% ± 2% | 95% ± 2% | 95% ± 2% | 90% ± 2% | 90% ± 2% | 91% ± 1% | 90% ± 2% | 94% ± 2% | |
| **Bag of worms** | | 4.5% ± 1% | 5% ± 2% | 5% ± 1% | 6.5% ± 1% | 7% ± 1% | 8.5% ± 1% | 9% ± 1% | 9% ± 2% | |
| **L1 Arrest** | | | | | 10% ± 1% | 10% ± 1% | 10% ± 1% | 25% ± 1% | 25% ± 1% | 45% ± 1% |
| **L3 Arrest** | | | | | 5% ± 1% | 5% ± 1% | 7% ± 1% | 10% ± 1% | 10% ± 1% | 20% ± 1% |
| **Slow or reduced movement** | + | + | + | + | + | + | + | ++ | ++ | ++ |
| *** This phenotype is visible from the larval stage 4 (L4).** | | | | | | | | | | |
| **** Fewer worms reach the L4 stage in F9 generation.** | | | | | | | | | | |

**Supplementary Table 1.** Phenotypes of continuously growing *xrn-2*(*PHX25*) worms at the non-permissive temperature of 26ºC.
